# Dual Role of a Rubisco Activase in Metabolic Repair and Carboxysome Organization

**DOI:** 10.1101/2020.05.16.099382

**Authors:** Mirkko Flecken, Huping Wang, Leonhard Popilka, F. Ulrich Hartl, Andreas Bracher, Manajit Hayer-Hartl

## Abstract

Rubisco, the key enzyme of CO_2_ fixation in photosynthesis, is prone to inactivation by inhibitory sugar phosphates. Inhibited Rubisco undergoes conformational repair by the hexameric AAA+ chaperone Rubisco activase (Rca) in a process that is not well understood. Here we performed a structural and mechanistic analysis of cyanobacterial Rca, a close homolog of plant Rca. In the Rca:Rubisco complex, Rca is positioned over the Rubisco catalytic site under repair and pulls the N-terminal tail of the large Rubisco subunit (RbcL) into the hexamer pore. Simultaneous displacement of the C-terminus of the adjacent RbcL opens the catalytic site for inhibitor release. An alternative interaction of Rca with Rubisco is mediated by C-terminal domains that resemble the small Rubisco subunit. These domains, together with the N-terminal AAA+ hexamer, ensure that Rca is packaged with Rubisco into carboxysomes. Cyanobacterial Rca is a dual-purpose protein with functions in Rubisco repair and carboxysome organization.

## INTRODUCTION

The AAA+ (ATPases associated with diverse cellular activities) chaperone Rubisco activase (Rca) serves as a paradigm of conformational enzyme repair (Bhat et al., 2017b; Mueller-Cajar, 2017). The exclusive client of Rca is the photosynthetic key enzyme Rubisco (ribulose-1,5-bisphosphate carboxylase/oxygenase), which is directly or indirectly responsible for all biomass production. Since Rca is required for optimal Rubisco function, understanding its mechanism is important in efforts to enhance photosynthesis with the goal to increasing crop yields (Andralojc et al., 2018; Bailey-Serres et al., 2019; Eva et al., 2019; Sharwood, 2017; Singer et al., 2019; Slattery and Ort, 2019).

Rubisco catalyzes the carboxylation of the 5-carbon sugar ribulose-1,5-bisphosphate (RuBP) in the Calvin-Benson-Basham cycle of photosynthesis. This multistep catalytic reaction produces two molecules of 3-phosphoglycerate as fuel for the synthesis of carbohydrates, fatty acids and amino acids. The most prevalent form of Rubisco (form I) in plants, algae and cyanobacteria is a hexadecamer of eight large (RbcL; ∼50-55 kDa) and eight small (RbcS, ∼12-18 kDa) subunits. The RbcL subunits form a cylindrical core, consisting of four anti-parallel dimers, capped by four RbcS subunits at each end of the cylinder. Two active sites per RbcL dimer unit are located at the interface between the N-terminal domain of one RbcL and the C-terminal domain of the other subunit. In order to become functionally competent, the RuBP binding site must be activated by carboxylation of the active-site lysine and binding of a Mg^2+^ ion, a process referred to as carbamylation. Upon binding of RuBP, the active site is sealed by a mobile loop from the C-terminal domain (the so-called loop 6) and by the otherwise flexible C-terminal sequence stretching over loop 6, forming a multilayered lid (Duff et al., 2000). Together with a loop sequence in the N-terminal domain of the adjacent subunit (the so-called ‘60s loop’), this generates the physical environment required for electrophilic attack of RuBP by CO_2_.

The requirement of Rca arises from the fact that the complex catalytic reaction of Rubisco is error prone, leading to the generation of tight-binding sugar phosphates that inactivate the enzyme. These “misfire” products include xylulose-1,5-bisphosphate (XuBP) and 2,3-pentodiulose-1,5-bisphosphate (PDBP) (Parry et al., 2008). Moreover, some plants under low light produce the inhibitory 2-carboxy-D-arabinitol-1-phosphate (CA1P) (referred to as “night-time” inhibitor). Binding of RuBP to non-carbamylated Rubisco also leads to inactive enzyme (Bracher et al., 2017; Parry et al., 2008; Portis et al., 2008). In all these conditions the release of the inhibitory sugar is mediated by Rca, which couples ATP hydrolysis to structural changes at the active site of Rubisco. Rca proteins have evolved independently in photosynthetic organisms and are associated with the red and green lineages of Rubisco (Bhat et al., 2017b; Mueller-Cajar, 2017; Tabita et al., 2008) (Figure 1A). While divergent in sequence, they all share the canonical AAA+ core consisting of an N-terminal α/β-nucleotide binding subdomain and a C-terminal α-helical subdomain (Figure 1A), and function as six-membered ring complexes. The AAA+ chaperones are typically substrate-promiscuous and act by threading flexible terminal sequences or loop segments of their target proteins into the hexamer pore (Olivares et al., 2016; Puchades et al., 2020). Threading is mediated by pore-loops (Figure 1A) that face the central pore and engage the substrate peptide, exerting a pulling force (Avellaneda et al., 2020; de la Pena et al., 2018; Dong et al., 2019; Fei et al., 2020; Ripstein et al., 2020; Rizo et al., 2019; Twomey et al., 2019).

**Figure 1.**
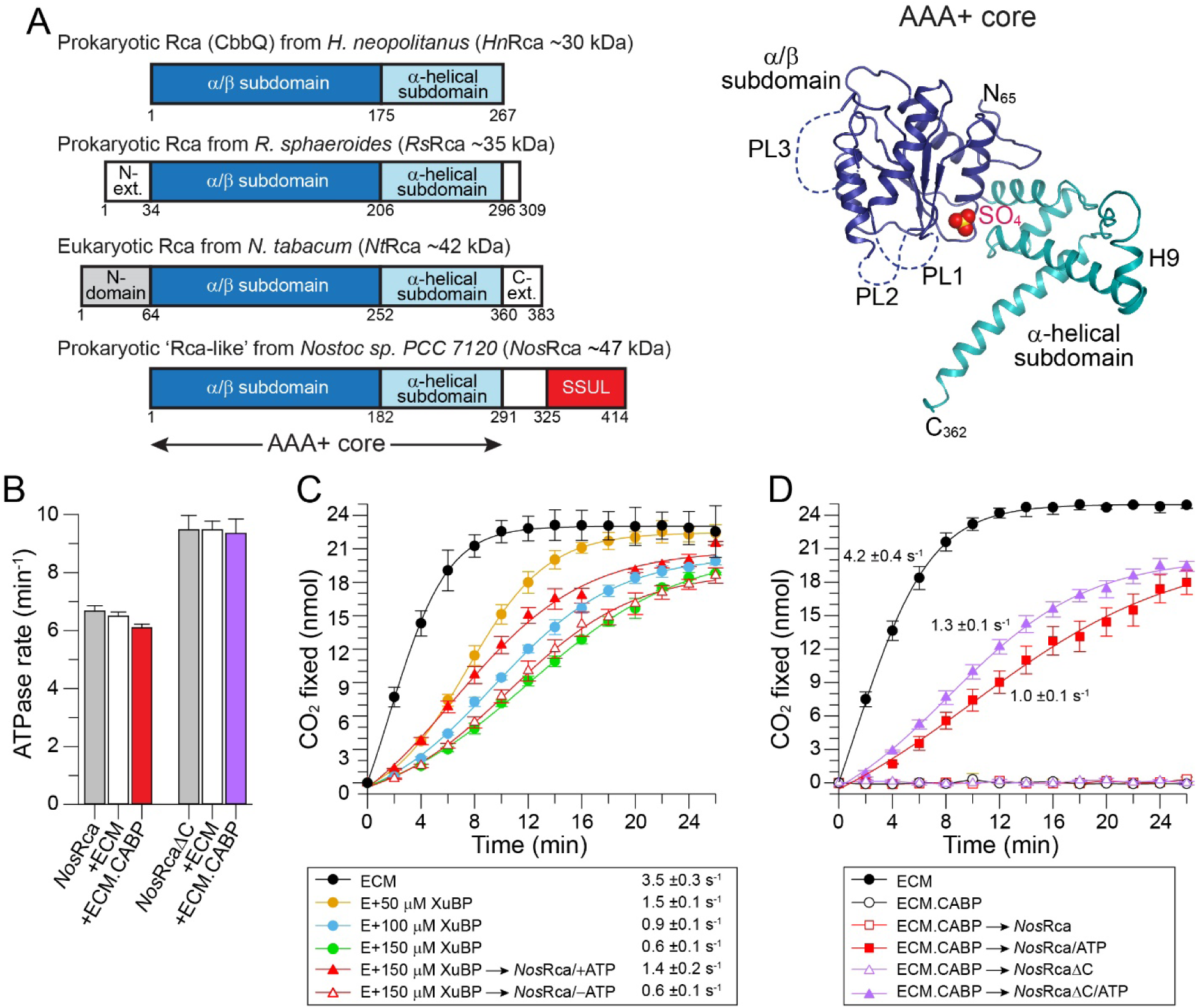
Rubisco Activase Function of *Nos*Rca. (A) Left: Schematic representation of the domain structures of Rca proteins across phylogenetic groups. CbbQ from the chemoautotroph *Halothiobacillus neapolitanus* (*Hn*Rca), CbbX from *R. sphaeroides* (*Rs*Rca; prokayotic, red-type), Rca from *N. tabacum* (*Nt*Rca; eukaryotic, green-type) and cyanobacterial ‘Rca-like’ (*Nos*Rca; prokaryotic, green-type) from *Nostoc* sp. PCC 7120. Right: Ribbon representation of Rca from *A. thaliana* (*At*Rca; PDB: 4W5W) (Hasse et al., 2015). α/β and α-helical subdomains as well as the pore-loops PL1, PL2 and PL3, and helix H9 are indicated. A bound sulfate ion bound to the nucleotide binding pocket is represented as space-filling model. (B) ATPase rates of *Nos*Rca and *Nos*RcaΔC in presence or absence of active (ECM) or inhibited (ECM.CABP) *Nos*Rubisco. ATPase rates were measured at 0.5 μM Rca (hexamer) and 0.25 μM NosRubisco (hexadecamer) as indicated. Bars represent averages with SD from at least three independent replicates. See STAR Methods for details. (C) *Nos*Rca functions as a Rubisco activase. CO_2_ fixation assays were performed with Rubisco enzyme (E, 0.25 μM) in presence of increasing concentrations of XuBP (50, 100 and 150 μM), and in presence or absence of 0.5 μM Rca (hexamer) and 3 mM ATP. The active *Nos*Rubisco (ECM) was used as control. Approximate rates of CO_2_ fixation were determined from the linear parts of the curves. Data represent SD from at least three independent replicates. (D) Reactivation of CABP inhibited *Nos*Rubisco (ECM.CABP) by *Nos*Rca or *Nos*RcaΔC. CO_2_ fixation assays were performed as in (C) with ECM.CABP in the presence or absence of ATP as indicated. Bars represent averages with SD of at least three independent replicates.

A low resolution cryo-electron microscopy (cryo-EM) structure of the Rca from the proteobacterium *Rhodobacter sphaeroides* (*Rs*) in complex with red-type Rubisco (form 1C) showed that the *Rs*Rca hexamer docks onto Rubisco over one catalytic site (Bhat et al., 2017a). *Rs*Rca is positioned so as to allow threading of the extended C-terminal sequence of *Rs*RbcL into the central pore, facilitating the opening of the catalytic site and release of the inhibitory sugar (Bhat et al., 2017a; Mueller-Cajar et al., 2011). In contrast, the mechanism of Rca in plants and green algae containing form IB Rubisco (Tabita et al., 2008) remains enigmatic, because form IB RbcL lacks the extended C-terminal sequence of red-type RbcL. Moreover, Rca of plants and green algae have a domain N-terminal to the AAA+ core that is required for Rubisco recognition (Figure 1A), and the so-called specificity helix H9 in the α-helical subdomain confers specificity for Rubisco proteins from solanaceous (nightshade) and non-solanaceous species (Li et al., 2005; Portis et al., 2008; Wachter et al., 2013). The mechanistic analysis of plant Rca has proven difficult, since the N-domain appears to be flexible and the protein, though active as a hexamer, populates a range of dynamic oligomeric states in vitro (Blayney et al., 2011; Keown and Pearce, 2014; Stotz et al., 2011).

The closest sequence homolog to plant Rca is the ‘Rca-like’ protein of several cyanobacteria with form IB Rubisco, which is required for elevated Rubisco activity under high light conditions (Li et al., 1993; Li et al., 1999). ‘Rca-like’ lacks the flexible N-terminal domain but its AAA+ core is highly homologous to eukaryotic Rca, with ∼38 % sequence identity and ∼70 % similarity (Data S1). A distinguishing feature is the presence of a Rubisco small subunit-like (SSUL) domain, connected to the C-terminus of the AAA+ core via a flexible linker (Lechno-Yossef et al., 2020) (Figure 1A). In cyanobacteria, Rubisco is packaged together with carbonic anhydrase into carboxysomes, membraneless compartments in which high concentrations of CO_2_ are generated for carbon fixation (Badger and Price, 2003; Price et al., 1998; Turmo et al., 2017). The scaffolding protein CcmM of β-carboxysomes utilizes multiple SSUL modules to induce Rubisco condensate formation during carboxysome biogenesis (Long et al., 2010; Wang et al., 2019). The SSUL domain of ‘Rca-like’ is highly homologous to these modules (∼40 % identity and ∼65 % similarity).

To obtain insight into the basic mechanism of plant Rca, here we analyzed the structure and function of the ‘Rca-like’ protein of the cyanobacterium *Nostoc* sp. PCC 7120 (*Nos*Rca). *Nos*Rca functions as a bona-fide Rubisco activase, independent of the SSUL domain. We solved the crystal structure of the stable *Nos*RcaΔC hexamer, and obtained its high resolution cryo-EM structure in complex with inhibited Rubisco. The structure of the complex revealed molecular details of the activation mechanism. A key finding is the well-resolved N-terminal amino acid sequence of RbcL inserted into the central pore of the spiral Rca hexamer. Additionally, we identified three interface regions, providing insight into how the Rca interacts with Rubisco and remodels the active site to liberate the inhibitory sugar phosphate. Mutational analysis confirmed that plant Rca also engages the N-terminal RbcL sequence of its cognate Rubisco, establishing N-terminal binding as the basic mechanism for activation of form IB Rubisco. We further show that the SSUL domains of the *Nos*Rca hexamer, by binding in a groove at the interface of RbcL dimer units, provide an alternative interaction resulting in network formation with Rubisco. The SSUL domains, together with the N-terminal AAA+ core which provides additional valency, ensure that Rca is packaged with Rubisco into carboxysomes. Cyanobacterial Rca thus combines two completely independent, and mutually non-exclusive functions in Rubisco remodeling (ATP-dependent) and carboxysome organization (ATP-independent).

## RESULTS

### *Nos*Rca Protein is a Rubisco Activase

The purified recombinant *Nos*Rca protein, with and without nucleotide, behaved as a ∼260 kDa complex by size-exclusion chromatography coupled to multiangle light scattering (SEC-MALS), consistent with a hexamer (theoretical mass ∼280 kDa) (Figures S1A and S1B, and Table S1). *Nos*Rca had a constitutive ATPase activity of ∼6.5 min^-1^ per protomer (Figure 1B). Since the AAA+ module shares a high degree of sequence identity (**∼**59 % identity and ∼88 % similarity) with the AAA+ modules of Rca from *Nicotiana tabacum* (*Nt*Rca) and *Arabidopsis thaliana* (*At*Rca), respectively (Data S2), we tested whether it functions as a Rubisco activase. Because activases are adapted to their cognate Rubisco (Bhat et al., 2017b; Mueller-Cajar, 2017), we recombinantly expressed and purified the Rubisco from *Nostoc* sp. PCC 7120 (*Nos*Rubisco) (Figure S1A) using coexpression of *Nostoc* chaperonin (*Nos*GroES/EL; see STAR Methods). The purified *Nos*Rubisco, activated with CO_2_ and Mg^2+^ (ECM), catalyzed the carboxylation of RuBP at a rate of ∼4 molecules of CO_2_ per active site s^-1^ (Figures 1C and 1D). However, as shown previously (Li et al., 1999), binding of RuBP to the non-activated enzyme did not result in inhibition (Figure S1C), in contrast to plant Rubisco (Parry et al., 2008; Stotz et al., 2011; Wang and Portis, 1992). We therefore tested the effect of the misfire sugar phosphate xylulose-1,5-bisphosphate (XuBP) as a competitive inhibitor (Bracher et al., 2015). A ∼60-80 % inhibition of the carboxylation rate was observed at concentrations of 50 to150 μ XuBP in the presence of 400 μ RuBP (Figure 1C), indicating that *Nos*Rubisco has a lower affinity for XuBP than plant Rubisco (Pearce, 2006). Addition of *Nos*Rca partially relieved the inhibition in an ATP-dependent manner (Figure 1C), suggesting that *Nos*Rca facilitated the release of XuBP from the Rubisco active sites. The effect was only partial, because the XuBP remained in the reaction as a competitive inhibitor. However, XuBP is unlikely to be a physiological inhibitor of *Nos*Rubisco, consistent with the absence in *Nostoc* sp. PCC 7120 of XuBP phosphatase, which serves to hydrolyze the XuBP upon release (Bracher et al., 2015). Complete and highly efficient inhibition of Rubisco was obtained with 2-carboxyarabinitol-1,5-diphosphate (CABP), a mimic of the carboxyketone intermediate of the carboxylation reaction, at a concentration equivalent to Rubisco active sites (0.25 μM) (Figure 1D). Strikingly, *Nos*Rca efficiently reactivated the inhibited Rubisco in the presence of ATP (Figures 1D). Thus, *Nos*Rca is a bona fide Rubisco activase, although the endogenous inhibitory sugar phosphate(s) of *Nos*Rubisco remains to be identified. Deletion of the C-terminal 123 residues (linker and the SSUL domain) in *Nos*Rca (*Nos*RcaΔC), resulted in a faster ATPase rate (∼9.5 min^-1^) and slightly more efficient Rubisco reactivation (Figures 1B and 1D), indicating that the SSUL domain is not required for activase function. The rate of reactivation was dependent on the concentration of *Nos*Rca (Figure S1D). Similar ATPase rates were measured in the presence of activated or inhibited Rubisco (Figure 1B), a property similar to plant Rca (Robinson and Portis, 1989).

### Crystal Structure of *Nos*RcaΔC

To allow a structural comparison of *Nos*Rca with plant Rca, we solved the crystal structure of *Nos*RcaΔC (residues 2-291) at ∼2.7 Å by Gd-multi-wavelength anomalous diffraction (Gd-MAD) (Figures S1A and Table S2). The resolution of the isomorphic native structure was 2.45 Å (Table S2). The asymmetric unit contained two subunits of *Nos*RcaΔC, with chain A (residues 2-275) having no nucleotide bound and chain B (residues 2-278) having ADP bound (Figures 2A, 2B and S2A). Note that the ADP bound during recombinant expression, as no nucleotide was added during purification and crystallization.

**Figure 2.**
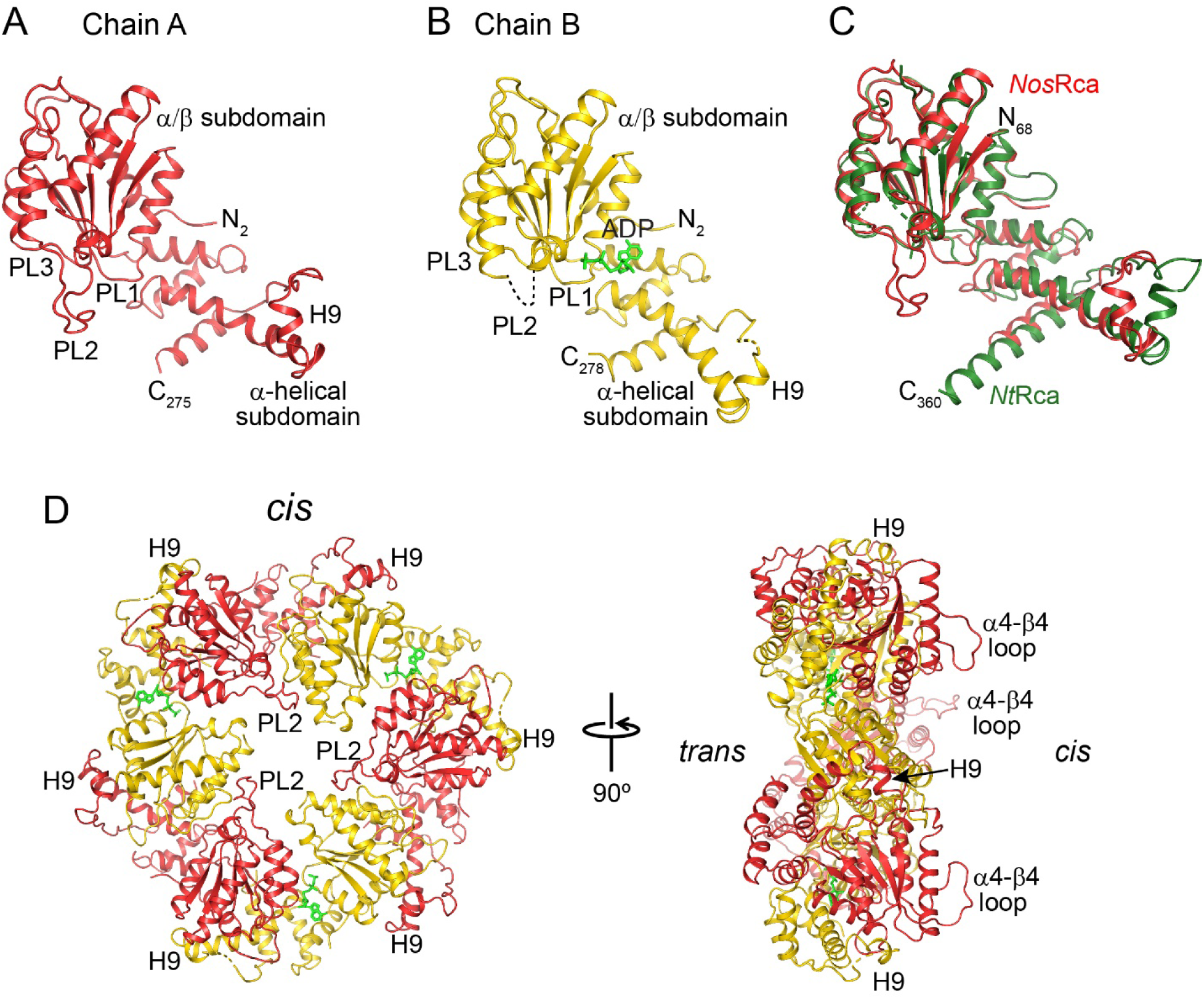
Crystal Structure of *Nos*RcaΔC. (A) and (B) Ribbon representations of the two crystallographically independent chains of *Nos*RcaΔC in red and yellow, respectively. The subdomain structure as well as locations of helix H9 and pore-loops PL1, PL2 and PL3 are indicated. ADP is represented as a wire-frame model in green. Chain termini are indicated. (C) Superposition of chain A (no nucleotide bound) of the asymmetric unit of *Nos*RcaΔC with *Nt*Rca (PDB: 3T15) (Stotz et al., 2011). Chain A of *Nos*RcaΔC and *Nt*Rca are shown as ribbons in red and green, respectively. (D) *Nos*RcaΔC hexamer. The hexamer was generated by applying the crystallographic three-fold rotational symmetry. The locations of helix H9 and pore-loop PL2 and loop α4-β4 are indicated. Chain A, red; chain B, yellow.

Overall the structure shows the typical AAA+ fold topology, consisting of the N-terminal Rossmann-fold α/β subdomain and a smaller C-terminal α-helical subdomain (Figures 2A and 2B). The two *Nos*RcaΔC subunits had nearly identical subdomain conformations, but differed in the inter-domain angle between the α/β and the α-helical subdomains by ∼11° (Figure S2B). Chain A superimposes well with the subunit structures of *Nt*Rca (r.m.s.d. 1.46 Å for 206 matching Cα positions) and *At*Rca (r.m.s.d. 1.96 Å for 219 matching Cα positions) (Figures 2C and S2C). The α/β-subdomains separately match almost perfectly (r.m.s.d. 0.53 Å and 0.52 Å, respectively). The structural conservation between *Nos*Rca and plant Rca includes the so-called specificity helix H9 (α9) in the α-helical subdomain (Figures 2A and 2B, and Data S1 and S2). As in the *Nt*Rca and *At*Rca crystal lattices (Hasse et al., 2015; Stotz et al., 2011), the α/β subdomain and helix H9 of the adjacent subunit form a rigid module (Figure S2D). The orientation of helix H9 to the four-helix bundle of the α-helical subdomain is flexible (Figure S2E). In the rhombohedral crystal lattice of *Nos*RcaΔC the subunits form a hexamer with alternating ADP-bound and nucleotide-free subunits (Figure 2D). Note that *Nos*RcaΔC is a hexamer in solution as analyzed by SEC-MALS, both in the absence and presence of added nucleotide (Table S1). The interface between the α/β subdomain and α-helical subdomain of adjacent subunits contains the nucleotide binding site (Figures 2D and S2A). In contrast to the pore-loops in the helical crystal structures of *Nt*Rca and *At*Rca (Hasse et al., 2015; Stotz et al., 2011), the pore-loops of *Nos*RcaΔC are ordered (PL1, residues 67-72; PL2, residues 105-119; PL3, residues 160-165), with the exception of the B chains where PL2 is disordered (Figures 2A, 2B and 2D). The hexamer pore would be wide enough for a sphere of ∼12 Å diameter to pass. In summary, the high degree of structural similarity of *Nos*Rca to plant Rca supports the identification of *Nos*Rca as a Rubisco activase. Thus, *Nos*Rca can serve as a useful model to understand the mechanism of plant Rca.

### *Nos*Rca Binds the N-terminus of RbcL in the Central Pore

To understand the mechanism of *Nos*Rca in remodeling the inhibited Rubisco, we sought to capture *Nos*Rca in the process of Rubisco reactivation by cryo-EM. We incubated CABP inhibited *Nos*Rubisco (ECM.CABP) with excess *Nos*RcaΔC in the presence of ATP, followed after 10 s by addition of the slowly hydrolyzing ATP analog, ATPγS (Figure 3A and STAR Methods), a nucleotide replacement strategy previously used in analyzing the 26S proteasome engaged with substrate (Dong et al., 2019). Cryo-EM analysis and single particle reconstruction yielded one predominant conformation of the *Nos*RcaΔC:Rubisco complex (Figures S3A-S3C). We obtained an EM density map of the complex with an overall resolution of ∼2.86 Å (Figures S3C, S3D and Table S3). The overall resolution of the Rubisco-subtracted *Nos*RcaΔC map was ∼3.29 Å (Figures S3C and S3E). *Nos*RcaΔC docks onto one corner of the cube-shaped Rubisco (Figure 3B), in a manner resembling the interaction of the prokaryotic red-type Rca from *R. sphaeroides* (Bhat et al., 2017a). The *cis* surface of the hexamer (Figure 2D) faces Rubisco with the N-terminal α/β-subdomains and the central pore of the hexamer being approximately positioned over one substrate binding pocket of a RbcL anti-parallel dimer unit (RbcL-A and RbcL-B; Figure 3B). The engaged catalytic site is in an open conformation with no discernable density for the inhibitor CABP, while the bound CABP was well-resolved in all other seven pockets of Rubisco (Figure S4A). Thus, the interaction between *Nos*RcaΔC and Rubisco captured in the cryo-EM structure represents the end-point of *Nos*Rca remodeling before dissociation from Rubisco.

**Figure 3.**
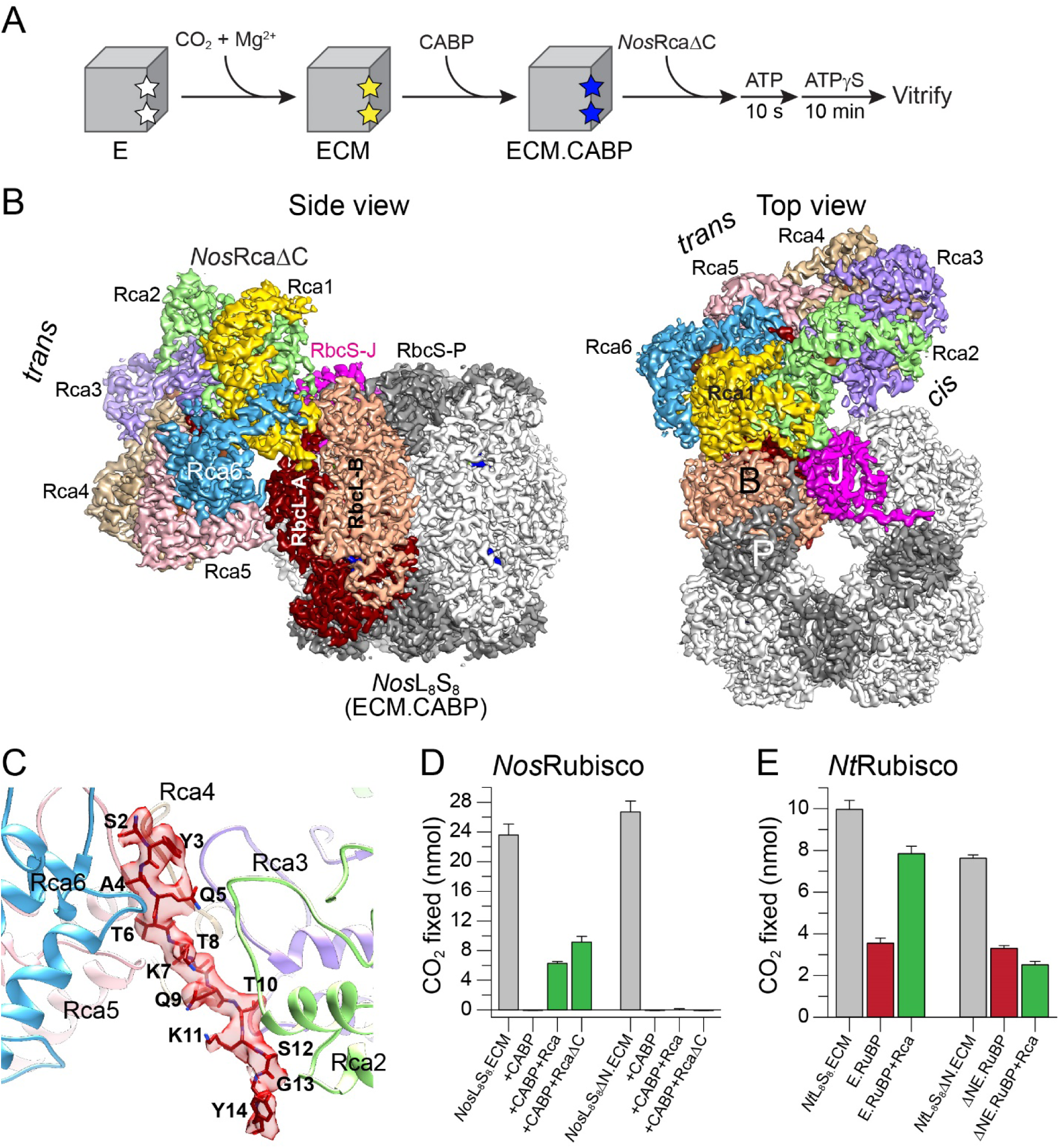
*Cryo-EM Structure of the Nos*RcaΔC:Rubisco Complex. (A) Schematic representation of preparation of the *Nos*RcaΔC:Rubisco complex analyzed by cryo-EM. (B) Cryo-EM reconstruction of the *Nos*RcaΔC:Rubisco complex in two perpendicular views. The subunits in the hexameric *Nos*Rca ring are labeled Rca1 to Rca6, with Rca1 (yellow) closest to and Rca6 (blue) farthest from Rubisco, respectively. The anti-parallel RbcL dimer subunits RbcL-A and RbcL-B as well as the RbcS subunits RbcS-J and RbcS-P in contact with NosRca are indicated. (C) Cryo-EM density of the N-terminal RbcL-A peptide bound inside the *Nos*Rca hexamer pore, interacting with Rca subunits (ribbon representation) Rca2 to Rca6. Amino acid residues of 2 to 14 of the bound peptide are indicated in single letter code. (D) Reactivation of *Nos*Rubisco by *Nos*Rca depends on the N-terminus of RbcL. CO_2_ fixation assays were performed as in Figure 1C with CABP-inhibited *Nos*Rubisco (*Nos*L_8_S_8_) and *Nos*L_8_S_8_ΔN in the presence or absence of *Nos*Rca or *Nos*RcaΔC and ATP for 8 min. Bars represent averages with SD from at least three independent experiments. (E) Reactivation of *Nt*Rubisco by *Nt*Rca depends on the N-terminus of RbcL. CO_2_ fixation assays were performed with RuBP-inhibited *Nt*L_8_S_8_ (E.RuBP) and *Nt*L_8_S_8_ΔN (ΔNE.RuBP) (0.25 μM active sites) in the presence or absence of *Nt*Rca (0.5 μM hexamer) and ATP for 8 min. Bars represent averages with SD from at least three independent experiments. See STAR Methods for details.

A key feature of the complex was the presence of well-defined density in the *Nos*RcaΔC hexamer pore, fitting to a 13-residue peptide in an extended conformation (Figure 3C). This finding suggested a mechanism similar to that of red-type proteobacterial Rca, which engages the flexible C-terminus of RbcL (Bhat et al., 2017a; Mueller-Cajar et al., 2011). However, the C-terminal sequence of RbcL-B was resolved up to residue Glu471 in the Rubisco structure, with only six residues (residues 472-477) missing in the EM density (Figures S4B and S4C), thus excluding engagement of the C-terminus by the *Nos*Rca central pore. Strikingly, we found that the N-terminal sequence of subunit RbcL-A residues Ser2 to Tyr14 fitted into the peptide density spanning the pore (Figure 3C). The next resolved N-terminal residue on RbcL-A was Thr24, with residues 15-23 connecting to the peptide in the pore being disordered (Figures S4B and S4D). In all other RbcL subunits the first ordered N-terminal residue is Gly13. We noted that the N-terminal sequence of *Nos*RbcL consists of alternating small and bulky side-chains (Figure 3C). Interestingly, this pattern is conserved at the N-terminus of green-type form IB RbcL (Tabita et al., 2008) of plants, green algae and other cyanobacteria (Figure S4E).

To test the importance of the N-terminus of RbcL for the reactivation reaction, we generated a truncated *Nos*Rubisco mutant lacking 12 residues at the N-terminus (*Nos*L_8_S_8_ΔN) (Figure S1A). *Nos*L_8_S_8_ΔN exhibited wild-type carboxylation activity, but could no longer be reactivated by *Nos*Rca upon inhibition with CABP (Figure 3D). To determine whether this mechanism is conserved in plants, we produced the analogous N-terminal RbcL truncation in Rubisco of *N. tabacum* (*Nt*L_8_S_8_ΔN) by recombinant expression in the presence of chloroplast chaperonins and assembly factors (Aigner et al., 2017; Lin et al., 2019; Wilson et al., 2019) (Figure S1A and STAR Methods). *Nt*L_8_S_8_ΔN was carboxylation active but the RuBP inhibited enzyme (E.RuBP) could not be reactivated by *Nt*Rca in contrast to wild-type Rubisco (Figures 3E and S1A, and STAR Methods). Thus, consistent with their high structural homology, engagement of the N-terminal RbcL sequence is important for remodeling by both *Nos*Rca and plant Rca.

### Pore-loop Interactions with the Rubisco N-terminal Peptide

The Rubisco-bound *Nos*RcaΔC hexamer adopted a helical “split-washer” conformation, with subunit 1 (Rca1; yellow) located closest to and Rca6 (blue) farthest from Rubisco (Figure 4A). Nucleotide density was present in all six nucleotide binding pockets of *Nos*Rca, ADP at the Rca1-Rca2 interface and ATP/ATPγS in the other sites (Figure S5A). The ATP/ATPγS was positioned close to the well-resolved side-chains of the arginine residues, Arg169 and/or Arg172, from the subsequent subunit (Figure S5A). These arginines have been implicated in ATP-hydrolysis in other AAA+ proteins, functioning as the so-called ‘arginine fingers’ (Puchades et al., 2020). The EM density of both arginines is less well defined at the Rca6-Rca1 interface, the split ends of the *Nos*Rca spiral (Figure S5A), perhaps reflecting the presence of mixed conformations due to on-going ATP hydrolysis.

**Figure 4.**
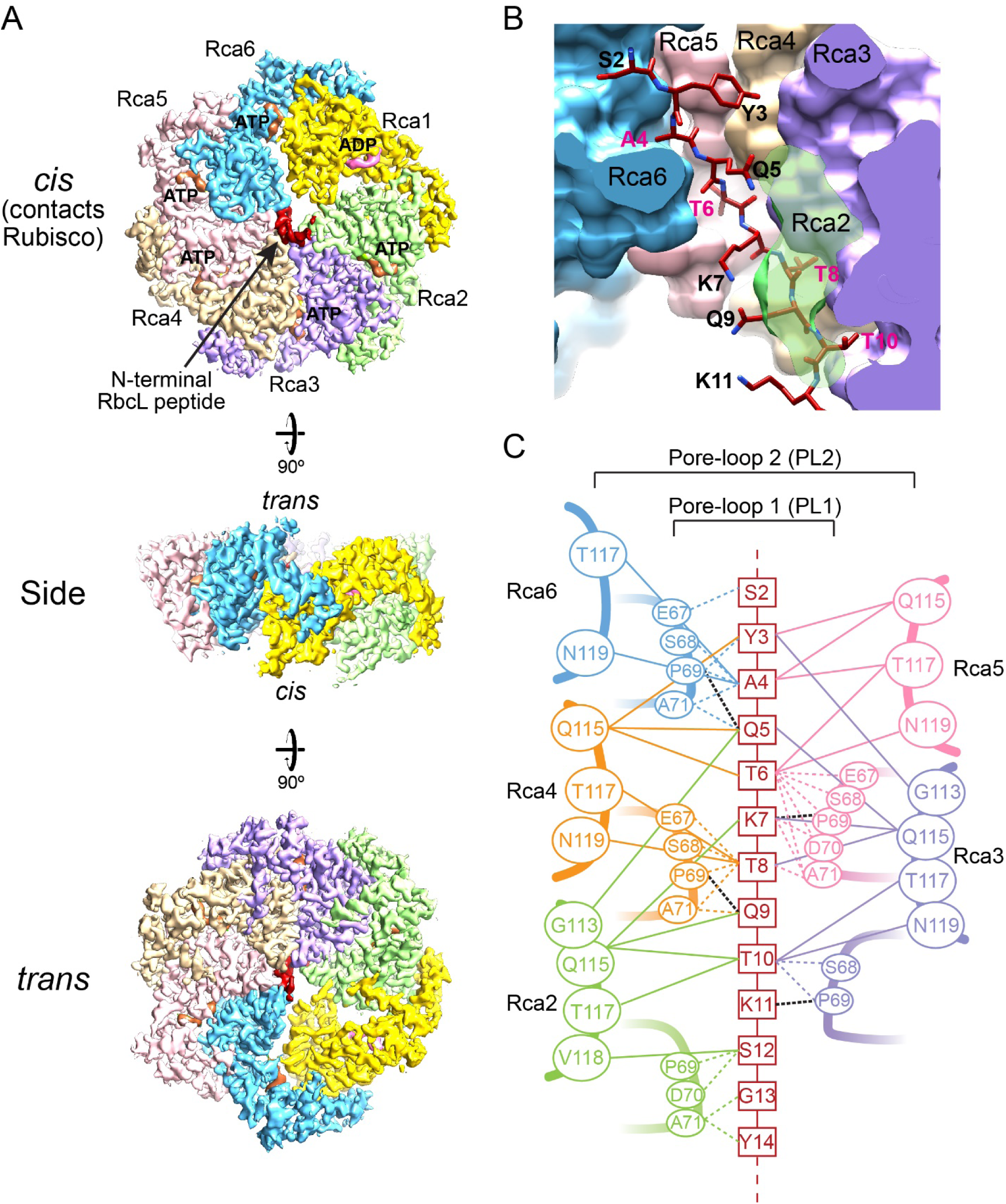
*Nos*Rca in a Helical ‘Split-washer’ Conformation Interacts via its Central Pore with the N-terminal RbcL Peptide. (A) Cryo-EM density map of *Nos*RcaΔC in the *Nos*RcaΔC:Rubisco complex in *cis* (surface in contact with Rubisco) and *trans* view. ADP in pink, ATP/ATPγS in sienna and the bound N-terminal peptide from RbcL-A in red. *Nos*Rca subunits are color coded as in Figure 3B. (B) Cavities in the *Nos*Rca central pore. Small cavities formed between pore-loops PL1 and PL2 from adjacent Rca subunits (Rca2 to Rca6) accommodate the side-chains of the alternating small amino acids of the RbcL N-terminal peptide. The bulky side-chains point to the solvent channel. The *Nos*Rca central pore is in surface representation and the RbcL N-terminal peptide in stick representation. (C) Interactions (van der Waals and hydrogen-bond) between residues of the RbcL N-terminal peptide and the staggered pore-loops PL1 (dotted colored lines) and PL2 (solid colored lines) of the Rca subunits. The distance cutoff was 4 Å. Putative hydrogen-bonds to the peptide backbone are indicated by black dotted lines.

The three pore-loops PL1 (residues 67-72), PL2 (residues 105-119) and PL3 (residues 160-165) (Data S2) were resolved in all *Nos*Rca subunits in the EM-density map (Figures S5B-S5D), adopting a staircase arrangement. Close examination of the bound peptide revealed that the small side-chains of RbcL-A residues Ala4, Thr6, Thr8 and Thr10 point into successive binding pockets in the central pore formed between the pore-loops PL1 and PL2 of adjacent subunits Rca2 to Rca6 (Figures 4B, S5B and S5C). The ADP-bound Rca1 is the only subunit not in contact with the peptide. The bulkier side-chains of Gln5, Lys7, Gln9 and Lys11 point into the pore solvent channel (Figure 4B). Note that Ser2 and Tyr3, protruding from the *trans* side of the pore, also conform to this pattern of alternating small and bulky side-chains, while the pattern does not continue beyond residue 12 (Figure S4E).

The bound peptide buries a substantial surface area of 1018 Å^2^ on *Nos*Rca. The contacts to the bound peptide are mediated by van der Waals interactions from both PL1 and PL2 (Figure 4C). In addition, the peptide backbone is in hydrogen-bond distance to the carbonyl group of Pro69 of PL1 in Rca3 to Rca6 (Figures 4C and S5E). Interestingly, the highly conserved aromatic residue Tyr116 in PL2, which is essential in plant Rca (Tyr188 in *Nt*Rca) (Stotz et al., 2011) (Data S2), does not make direct contact with the peptide (Figure S5F). In the ATP-bound subunits Rca2 to Rca6, Tyr116 instead forms a hydrogen-bond with Gln121 in the subsequent subunit, resulting in a network of interactions that rigidifies the central pore (Figure S5F). In contrast, the position of Tyr116 is ill-defined in the ADP-bound Rca1. PL3, which makes no contact with the bound peptide (Figure S5D), is positioned between the nucleotide binding site and PL2, suggesting that it may be involved in allosteric regulation via the spatially adjacent arginines Arg169 and Arg172.

### Details of the *Nos*RcaΔC:Rubisco Interface and Remodeling Mechanism

Beyond engaging the N-terminal sequence of RbcL, *Nos*Rca contacts Rubisco at three regions covering a total of 1445 Å^2^ (Figures 5A-5E). Interface I (419 Å^2^) is formed between Rca5/Rca6 and RbcL-A, and has high surface shape complementarity (Figures 5A and 5B). Rca5 residues Leu244 and Asn248 of the specificity helix H9 as well as Leu250, and Rca6 residues Val91 and Gly93 make van der Waals contacts with Glu52, Ile88, Pro90, Pro92 and Asp95 of RbcL-A (Figure 5B). Arg92 in helix α3 of Rca6 forms a salt bridge with Glu52 of RbcL-A, and an intramolecular van der Waals interaction with Glu57 (Figure 5B). Interfaces II and III involve the α4-β4 loop of Rca1 and Rca2, respectively (Figures 5C-5E). In interface II (600 Å^2^), subunit Rca1 inserts between the RbcL-A and RbcL-B antiparallel dimer (Figure 5C). The conformation of the protruding α4-β4 loop is rigidified by an intramolecular salt bridge formed between Arg83 and Asp144, with Pro140 having a further stabilizing effect on the loop. The α4-β4 loop contacts Trp67 of the substrate binding pocket, which is located in the 60s loop (residues 60-69) of RbcL-A. In addition, Tyr143 of Rca1 makes van der Waals contacts with Gln46, Lys129 and Ala130 of RbcL-A (Figure 5C). Note that there is also a weak interaction between Val91 of Rca1 and the C-terminal residues Ile466 and Lys467 of RbcL-B, which might help in keeping the substrate pocket open for unhindered inhibitor release (Figure 5C). In interface III (431 Å^2^) the α4-β4 loop of Rca2 contacts the RbcS subunits J and P (Figures 5D and 5E). Specifically, Tyr143 of Rca2 engage in van der Waals interactions with the β1-α2 loop (K57-L61) of RbcS-J and Met1 of RbcS-P (Figure 5E). Residues Arg92, Gly93 and Asn248 of interface I, as well as Pro140 and Tyr143 of interface II/III are conserved in plant Rca (Data S1), providing additional support for a conserved mechanism of remodeling of green-type form IB Rubisco.

**Figure 5.**
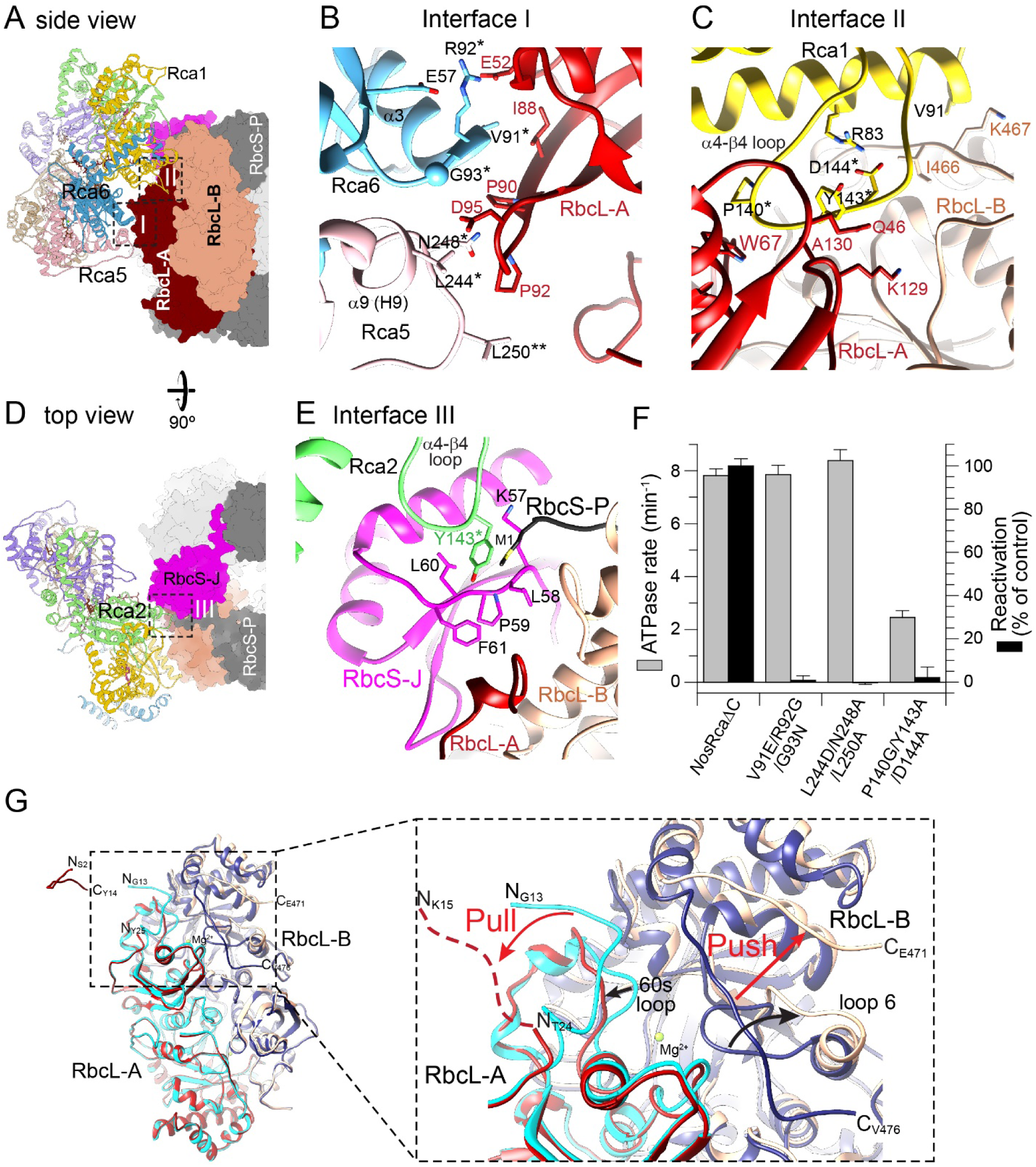
Structural Details of the *Nos*Rca:Rubisco Interaction. (A-E) Interfaces of the *Nos*RcaΔC:Rubisco complex. Overview of the complex, highlighting interfaces I and II with *Nos*Rca in ribbon representation and Rubisco in surface representation (A). Interacting amino acid residues in interface I (B) and interface II (C). In interface I, Rca5 and Rca6 (subunits color-coded as in Figure 3B) interact with RbcL-A. Helix α9 (H9) in Rca5 and helix α3 in Rca6 are indicated. In interface II, the α4-β4 loop of Rca1 makes contacts with RbcL-A. Overview of the complex highlighting interface III (D). In interface III, the residue Tyr143 of the α4-β4 loop of Rca2 makes contacts with RbcS-J and RbcS-P (E). Sidechains of contact residues are shown in stick representation. *, residue analyzed by mutation. (F) Mutational analysis of *Nos*Rca interface residues. ATPase rates of *Nos*RcaΔC and mutants (*Nos*RcaΔC V91E/R92G/G93N, *Nos*RcaΔC P140G/Y143A/D14A, *Nos*RcaΔC L244D/N248A/L250A) and reactivation of CABP-inhibited *Nos*Rubisco were measured as in Figures 1B and 3D, respectively. ATPase rates were measured in the absence of Rubisco at 20 mM KCl (see STAR Methods). CO_2_ fixation was measured for 8 min and set to 100 % for wild-type *Nos*RcaΔC. Error bars represent SD of at least three independent experiments. (G) Structural remodeling of the Rubisco substrate binding pocket. Super-position of the binding pocket, formed at the interface between the N- and C-terminal domains of the anti-parallel RbcL subunits, in the closed (RbcL-A light blue; RbcL-B dark blue) and the *Nos*Rca-engaged open state (RbcL-A red; RbcL-B peach). The remodeled regions, including the 60s loop of RbcL-A and loop 6 of RbcL-B as well as the pull and push actions of *Nos*Rca are indicated.

Next, we performed a mutational analysis of *Nos*RcaΔC to test the functional significance of these interactions. Note that the integrity of the hexamer structure was preserved in the mutant proteins (Table S1). The interface I mutants V91E/R92G/G93N (breaking the Arg92-Glu52 salt bridge and disrupting the van der Waals interactions with Ile88 and Pro90) and L244D/N248A/L250A (disrupting the interaction with Asp95 and Pro92) preserved wild-type ATPase activity, but failed to reactivate the CABP-inhibited Rubisco (Figures 5F and S1A). The interface II/III mutant P140G/Y143A/D144A (disrupting the interaction of the α4-β4 loop with Rubisco and destabilizing the loop) also abolished reactivation, but additionally showed a ∼70 % reduced ATPase activity (Figures 5F and S1A). Mutational analyses in various plant Rca proteins have also implicated residues in helix α3, helix α9 and the α4-β4 loop in the interaction with Rubisco (Li et al., 2005; Ott et al., 2000; Portis et al., 2008; Shivhare and Mueller-Cajar, 2017; Shivhare et al., 2019), supporting the functional significance of the structurally defined interface regions in the *Nos*RcaΔC:Rubisco complex.

How does *<u>Nos</u>*Rca remodel Rubisco? The *Nos*RcaΔC:Rubisco complex represents a post-remodeling state, as the Rubisco substrate binding pocket that is engaged by Rca is in an open conformation with no inhibitor bound (Figure S4A). Super-position of the open and closed conformations of the substrate binding pockets revealed the structural changes effected by *Nos*Rca (Figure 5G): i) The N-terminal residues 15-23 of RbcL-A are disordered, with no clear discernable EM density, but are well resolved in all other subunits, suggesting that binding the N-terminal sequence (residues 2-14) in the central Rca pore substantially destabilizes this region; ii) As a consequence, the 60s loop of RbcL-A, which contributes to stabilizing the substrate binding pocket, is displaced (Figure 5G); iii) In addition, the Rca hexamer – with the bulk of Rca1 acting as a wedge – displaces the C-terminal peptide of the adjacent RbcL subunit from its closed position over the substrate pocket (RbcL-B), thereby disrupting its interaction with loop 6. As a result, loop 6 is retracted from its closed position over the substrate binding pocket (Figure 5G). The combined displacement of the 60s loop and loop 6, both of which point to the solvent in the post-remodeling state, results in the complete opening and destabilization of the substrate binding site, facilitating inhibitor release.

### Function of the SSUL Domains in Carboxysome Organization

A distinguishing feature of the *Nos*Rca subunit is the presence of the C-terminal SSUL domain (Figure 1A), which is connected to the AAA+ core (residues 2-291) by a ∼35 residue flexible linker. The last structured residue in the EM density map of all *Nos*RcaΔC subunits is Asp281. The SSUL domain shares 54-64 % sequence identity with the three SSUL modules of the CcmM protein (*Nos*M58) and its shorter variant M35 (*Nos*M35) (Long et al., 2010) (Figure 6A and S6A). The SSUL modules of CcmM function as scaffolding proteins to concentrate Rubisco into a dense condensate prior to encapsulation into the β-carboxysome (Ryan et al., 2019; Wang et al., 2019). Thus, it seemed plausible that the SSUL domains recruit *Nos*Rca into the Rubisco condensate (Lechno-Yossef et al., 2020). Notably, while in M58 and M35 three to five SSUL modules are linked to provide the multi-valency for Rubisco network formation, in *Nos*Rca each subunit of the hexamer carries its own SSUL, presumably generating high avidity.

**Figure 6.**
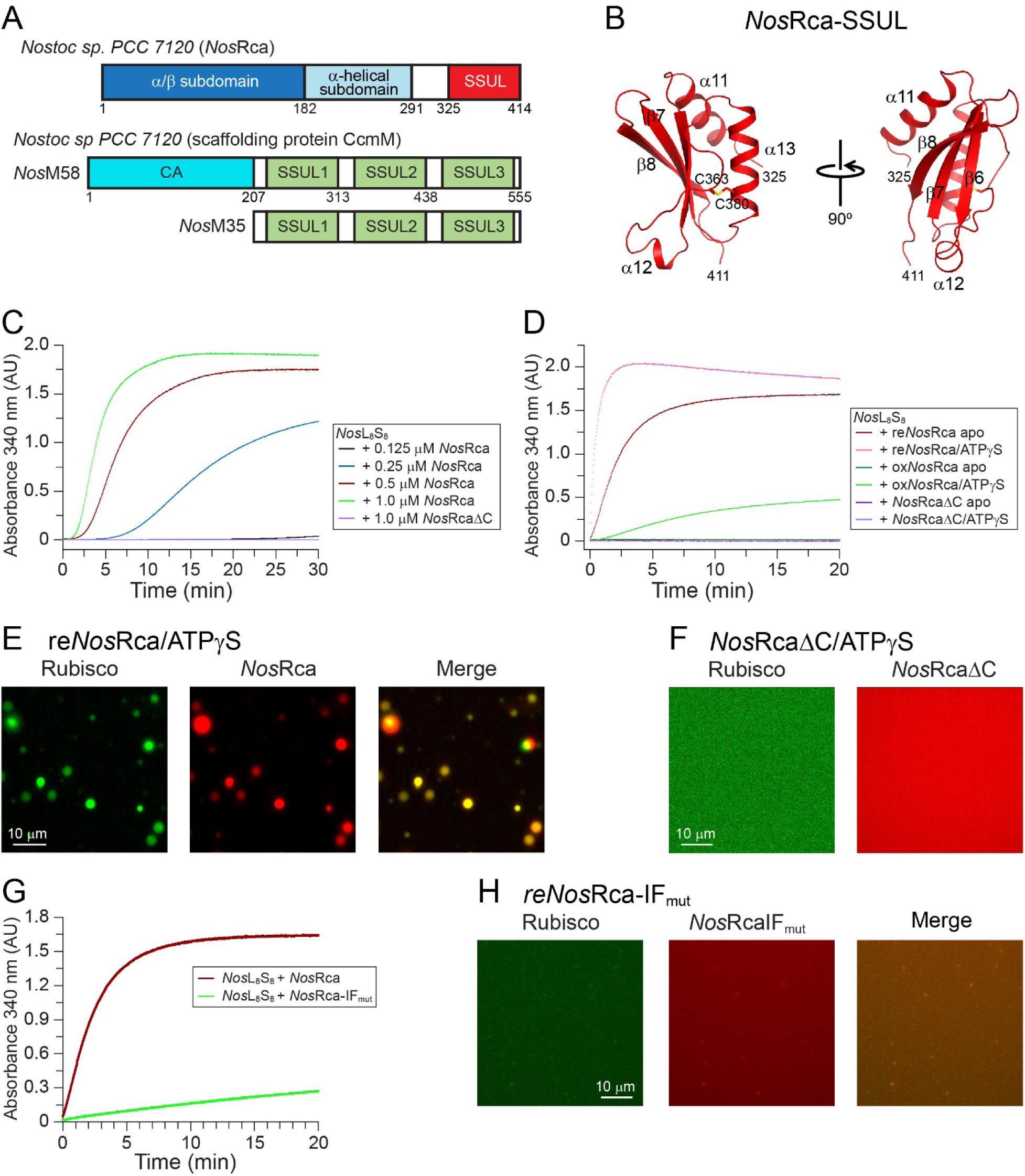
Cooperation of SSUL Domains and AAA+ Core in Carboxysome Organization. (A) Schematic representation of the domain structures of *Nos*Rca and *Nos*CcmM (full length *Nos*M58 and the shorter isoform *Nos*M35). CA, carbonic anhydrase domain. (B) Crystal structure of the SSUL domain of *Nos*Rca in ribbon representation. Chain termini, secondary structure elements and the disulfide bond (Cys360-Cys380) are indicated. (C) Formation of *Nos*Rca-Rubisco condensate mediated by the SSUL domains analyzed by turbidity assay. Data shown are for Rubisco (*Nos*L_8_S_8_; 0.25 μM) with increasing concentrations of reduced *Nos*Rca from 0.125 μM to 1 μM or with *Nos*RcaΔC (1 μM). Representative results from at least three independent experiments are shown. (D) Nucleotide and redox-dependence of *Nos*Rca-Rubisco condensate formation. Rubisco (0.25 μM) was combined with reduced (re) or oxidized (ox) *Nos*Rca (0.5 μM), or *Nos*RcaΔC (0.5 μM), in the presence or absence of ATPγS (2 mM). Condensate formation was measured by turbidity assay as in (C). Representative results from at least three independent experiments are shown. (E and F) Formation of *Nos*Rca-Rubisco condensate in the presence of ATPγS as observed by fluorescence microscopy. Fluorophore labeled proteins (A405-*Nos*Rca, A405-*Nos*RcaΔC and A532-*Nos*Rubisco) were mixed 1:10 with the respective unlabeled proteins and used at total concentrations of reduced *Nos*Rca and Rubisco of 0.5 μM and 0.25 μM, respectively. Rubisco was mixed with either reduced *Nos*Rca (E) or *Nos*RcaΔC (F) in the presence of 2 mM ATPγS. See STAR Methods for details. (G) *Nos*Rca-Rubisco condensate formation analyzed as in (D). Rubisco (0.25 μM) was combined with reduced *Nos*Rca or *Nos*Rca-IF_mut_ (0.5 μM). Representative results from at least three independent experiments are shown. (H) Inability of *Nos*Rca-IF_mut_ to form a condensate with Rubisco, observed by fluorescence microscopy. Fluorophore labeled A405-*Nos*Rca-IF_mut_ and A532-*Nos*Rubisco were used as described in (E).

The crystal structure of *Nos*Rca*-*SSUL (*Nos*SSUL, residues 325-414; Figure 1A), solved by MAD at 1.4 Å resolution (Figures 6B, S6B and S6C, and Table S2), confirmed the high degree of similarity to the structure of the SSUL1 module in CcmM of *Synechococcus elongatus* PCC 7942 (*Se*SSUL) (r.m.s.d. 1.02–1.04 Å for matching Cα positions) (Figure S6D). Like *Se*SSUL, *Nos*SSUL contains a disulfide bond (Cys363 to Cys380) in the crystal (Figure 6B). The redox state of *Se*SSUL has been shown to regulate Rubisco condensate formation (Wang et al., 2019), with the cytosol being reducing and the carboxysome interior oxidizing (Chen et al., 2013; Pena et al., 2010; Price et al., 1992).

To investigate whether *Nos*Rca has the ability to induce Rubisco condensate formation similar to *Se*CcmM, we performed an established turbidity assay (Wang et al., 2019). We incubated Rubisco (0.25 μM hexadecamer) with increasing concentrations of reduced *Nos*Rca (0.125 to 1.0 μM hexamer). Turbidity at 340 nm developed within minutes in a manner dependent on *Nos*Rca concentration (Figure 6C), consistent with a recent report for the ‘Rca-like’ protein from *Fremyella diplosiphon* (Lechno-Yossef et al., 2020). No turbidity was observed with *Nos*RcaΔC (Figure 6C), indicating that the interaction with Rubisco is mediated by the SSUL domains. Neither was turbidity observed with oxidized *Nos*Rca (0.5 μM), indicating that the presence of the disulfide bond in SSUL reduces the affinity for Rubisco (Wang et al., 2019) (Figure 6D and STAR Methods). Note that the oxidized *Nos*Rca was as ATPase active and functional in Rubisco remodeling as the reduced protein (Figure S6E). Cryo-EM analysis resolved the interaction of *Nos*Rca with Rubisco mediated by *Nos*SSUL at 8.3 Å resolution (Figures S7A-S7D). Docking of the *Nos*SSUL crystal structure and the *Nos*Rubisco model into the low resolution EM density map revealed that the SSUL module binds close to the equatorial region of the hexadecamer in a groove between RbcL dimers (Figure S7E), as previously shown for the *Se*M35-Rubisco complex (Wang et al., 2019).

Interestingly, addition of ATPγS to reduced *Nos*Rca resulted in enhanced turbidity with Rubisco (Figure 6D). However, this effect was still completely dependent on the presence of the SSUL domains (Figure 6D), suggesting the possibility that the AAA+ core modulates the interaction by contributing additional valency. Consistent with such an enhancing effect, addition of ATPγS also enabled measurable turbidity with oxidized *Nos*Rca (Figure 6D).

To determine whether the observation of turbidity reflected demixing of *Nos*Rca and Rubisco into phase separated droplets, we N-terminally labeled Rubisco with Alexa Fluor 532 (Rubisco-A532) and *Nos*Rca (or *Nos*RcaΔC) with Alexa Fluor 405 (A405-Rca or A405-RcaΔC). Upon incubation of the Rca protein with Rubisco (labeled:unlabeled 1:10), liquid-liquid phase separation (LLPS) into fluorescent droplets was readily observed (Figure 6E). No droplet formation occurred with *Nos*RcaΔC and Rubisco (Figure 6F).

To directly assess the contribution of the AAA+ core to LLPS, we generated a mutant of *Nos*Rca with disrupted interface I (mutations V91E/R92G/G93N and L244D/N248A/L250A combined) (*Nos*Rca-IF_mut_). *Nos*Rca-IF_mut_ was a stable hexamer in solution (Figure S1A and Table S1). Strikingly, the ability of *Nos*Rca-IF_mut_ to induce Rubisco condensate formation was strongly reduced (Figure 6G), and only very small droplets were observed, despite the presence of the SSUL domains (Figure 6H). Thus, in addition to the SSUL domains, the interaction via the AAA+ core contributes critical valency to the phase separation process. Apparently, both interaction modes cooperate in ensuring packaging of *Nos*Rca into carboxysomes together with Rubisco.

## DISCUSSION

### Mechanism of Form 1B Rubisco Metabolic Repair

Our structural and functional analysis of cyanobacterial *Nos*Rca, a close homolog of plant Rca, revealed the mechanism of ATP-dependent form 1B Rubisco metabolic repair. In the high-resolution cryo-EM structure of the *Nos*RcaΔC:Rubisco complex, *Nos*Rca docks onto inhibitor-bound Rubisco with its hexamer pore approximately positioned over one of the eight catalytic sites. In contrast to the proteobacterial red-type *Rs*Rca, which engages the extended RbcL C-terminus (Bhat et al., 2017a), *Nos*Rca is rotated clockwise by ∼50° degrees, positioning its central pore to bind the N-terminus of RbcL (Figure 7A). Insertion of the N-terminus into the central pore of *Nos*Rca is critical for Rubisco activation. This mechanism is likely of general relevance in organisms with green-type form 1B Rubisco (cyanobacteria, green algae and plants), as their RbcL subunits lack the extended C-terminal sequence of red-type RbcL. Indeed, N-terminal truncation of RbcL from tobacco prevented activation of inhibited Rubisco by its cognate Rca.

**Figure 7.**
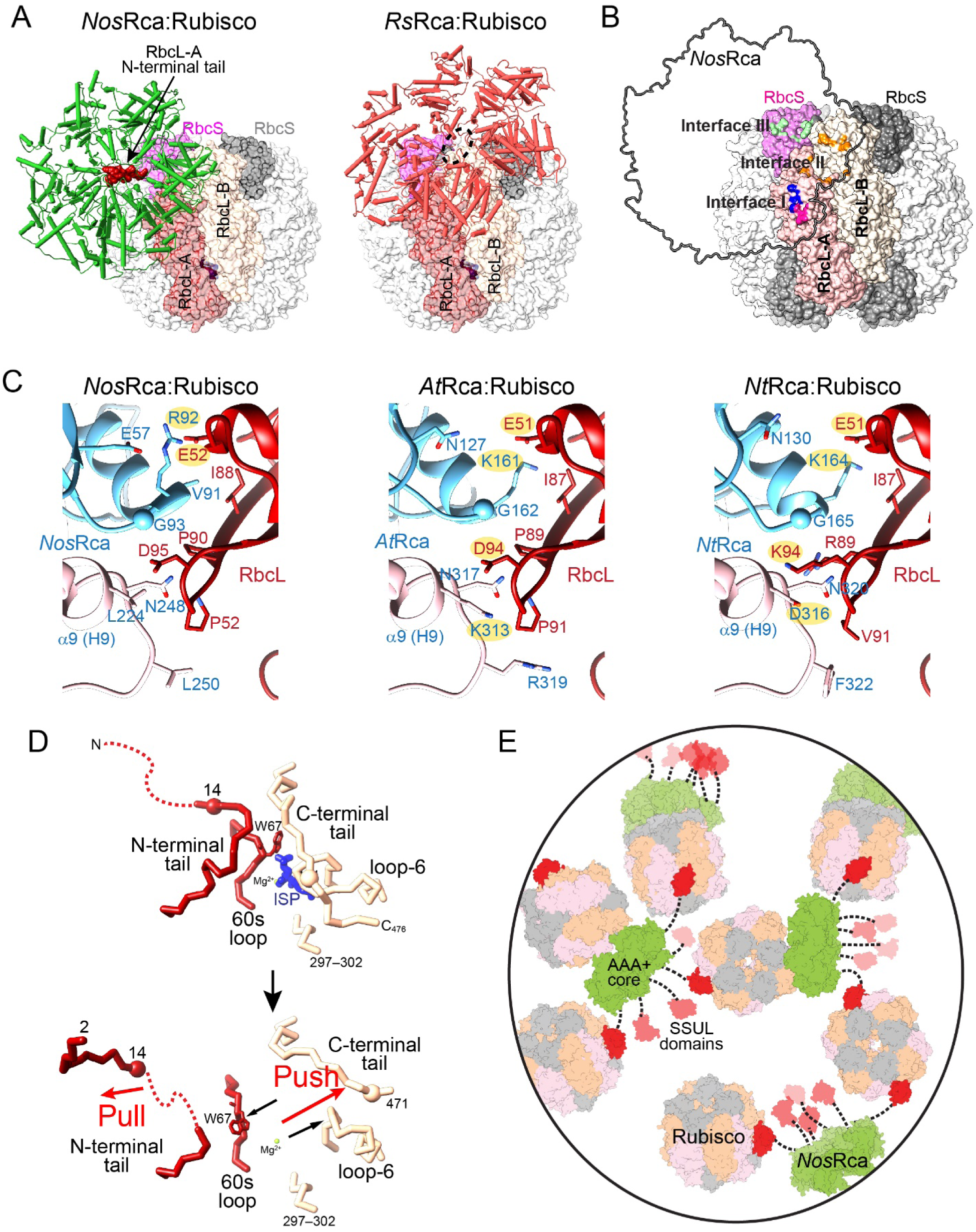
Mechanism of *Nos*Rca in Rubisco Reactivation and Carboxysome Organization. (A) Comparison of the positioning of *Nos*Rca (left) and *Rs*Rca (right) in complex with their cognate Rubisco for engagement of the N-terminus and C-terminus of RbcL, respectively. In both structures Rubisco is shown in surface representation in the same orientation.The anti-parallel RbcL dimer and the RbcS subunits involved in the interaction with Rca are indicated. *Rs*Rca:Rubisco model, EMDB: EMD-3701/PDB: 3ZUH and PDB: 5NV3. (B) Overview of the surface areas on *Nos*Rubisco interacting with *Nos*Rca via interfaces I (red and dark blue), II (orange) and III (light blue). *Nos*Rca is shown as an outline. *Nos*Rubisco colors as in (A). (C) Structural models for interface I in the *At*Rca:*At*Rubisco (middle) and *Nt*Rca:*Nt*Rubisco (right) complexes. On the left, interface I in the *Nos*Rca:*Nos*Rubisco complex is shown. The protein backbones are shown in ribbon representation. Interface sidechains are shown in stick representation. The hypothetical models were created by superposing the α-helical subdomains from crystal structures of *At*Rca (PDB: 4W5W) (Hasse et al., 2015) and *Nt*Rca (PDB: 3T15) (Stotz et al., 2011) onto *Nos*Rca. Sequence differences to *Nostoc* and sidechain conformers were manually adjusted. (D) Structural rearrangement of the Rubisco substrate binding pocket by *Nos*Rca from the closed state with inhibitory sugar phosphate (ISP) bound to the open state (ISP released). (E) Cooperation of the SSUL domains and the AAA+ core of *Nos*Rca in Rubisco interaction during carboxysome biogenesis. The model is based on the interactions observed in the cryo-EM reconstructions of *Nos*RcaΔC:Rubisco and *Nos*Rca:Rubisco (showing *Nos*SSUL-Rubisco interaction) in the absence of nucleotide. The dotted lines represent the ∼35 residue flexible linker between the AAA+ core (green) and the SSUL domains (red).

The complex of *Nos*Rca with inhibitor-bound Rubisco, stabilized by a nucleotide substitution strategy, represents an end-state of the remodeling process in which the substrate binding site under repair is open and the inhibitor released. In addition to insertion of the N-terminus of one RbcL (RbcL-A) into the Rca hexamer pore, destabilization of the binding pocket involves extensive interactions with Rubisco via three interface regions (Figure 7B). Underscoring the sequence homology to plant Rca (Data S1), critical residues of these interfaces are structurally conserved and their functional relevance is confirmed by mutagenesis (Figure 7C). Interface I formed by Rca5/Rca6 with RbcL-A includes the specificity helix H9 on Rca5 (Figure 5B). In plants, helix H9 functions in distinguishing Rubisco proteins from solanaceae and non-solanaceae (Portis et al., 2008; Wachter et al., 2013). Notably, the RbcL residues Pro90 and Asp95, which mediate Rca specificity in non-solanaceous plants (e.g. Pro89 and Asp94 in *A. thaliana* and *Spinacia oleraces*), are conserved in *Nos*RbcL (Li et al., 2005; Ott et al., 2000) (Figure 7C). Pro90 contacts the *Nos*Rca backbone at Val91 and Gly93 in the α3-β3 loop of Rca6. Note that in plant Rca sequences this loop is one residue shorter (Data S1 and S2). Asp95 in *Nos*RbcL makes van der Waals contacts to Leu244 in helix H9 of *Nos*Rca5 (Figure 7C). Modeling of the *At*Rca:Rubisco complex based on the *Nos*Rca:Rubisco structure suggests that Asp94 in *At*RbcL is in electrostatic interaction with Lys313 in *At*Rca (the residue equivalent to Leu244 in NosRca) (Figure 7C). This interaction mediates the Rca-Rubisco specificity of non-solanaceous plants, consistent with previous mutational analyses (Li et al., 2005; Ott et al., 2000; Portis et al., 2008). In tobacco, a solanaceous plant, this electrostatic interaction is reversed, with Lys94 on *Nt*RbcL interacting with Asp316 in *Nt*Rca (Figure 7C), and the charge reversal switching the specificity from non-solanaceous to solanaceous (Portis et al., 2008). Another conserved interaction in interface I is the salt bridge Arg92 in *Nos*Rca6 with Glu52 in RbcL, which is equivalent to the salt bridges of Lys161 to Glu51 and Lys164 to Glu51 in *A. thaliana* and tobacco, respectively (Figure 7C).

Interfaces II and III involve the α4-β4 loop of Rca1 and Rca2. In interface II, the α4-β4 loop of Rca1 inserts between the RbcL-A and RbcL-B subunits of the antiparallel dimer containing the catalytic site under repair. In interface III, the α4-β4 loop of Rca2 makes contact with RbcS subunits J and P. As a result of these combined interactions, the catalytic site is converted to a completely open state. Specifically, the 60s loop of RbcL-A, and the C-terminal tail and loop 6 of the adjacent RbcL-B subunit, all of which contribute to stabilizing the closed state of the substrate binding pocket, are displaced (Figure 7D). The interaction of *Nos*Rca with RbcS is intriguing. RbcS contributes indirectly to stabilizing the sugar substrate in the active site pocket (Bracher et al., 2011) and thus the interaction with *Nos*Rca, in addition to providing an anchoring point, may also prevent dissociation of RbcS while the catalytic site is being destabilized.

### Binding of the RbcL N-terminus in the Rca Central Pore

The N-terminal peptide of RbcL-A (residues 2-14) buries an extensive surface of the *Nos*Rca pore, apparently representing an end-state in which Rca spends most of its time during the remodeling cycle. Consistent with this assumption, the *Nos*Rca:Rubisco complex was resolved only in one defined state. Presumably, ATP-dependent peptide threading comes to a halt when this stably locked position is reached, thereby limiting further Rca action.

The interactions of the N-terminal residues of RbcL-A with the central Rca pore are remarkably well resolved in the cryo-EM structure. This sequence is characterized by an alternating pattern of small and bulky residues, which is generally conserved in green-type form 1B RbcL. The small side-chains of Ala4, Thr6, Thr8 and Thr10 are pointing into successive binding pockets in the central pore, while Ser2 and Tyr3 protrude from the *trans*-side of the pore. The binding pockets appear to represent an adaptation to the alternating side-chain pattern of the RbcL N-terminus. They are formed by pore-loops PL1 and PL2 of adjacent subunits from Rca2 through Rca6, and are staggered due to the stair-case arrangement of the Rca subunits. In contrast, the extended C-terminal sequence of red-type RbcL does not show a conserved side-chain pattern (Mueller-Cajar et al., 2011). Structural information on the interaction of the red-type RbcL C-terminus with the central pore of its cognate Rca is still missing.

Many AAA^+^ proteins, including *Rs*Rca, contain an aromatic residue (Tyr114 in *Rs*Rca) in their canonical PL1 motif (aromatic-hydrophobic-glycine) that is involved in substrate peptide threading (de la Pena et al., 2018; Dong et al., 2019; Fei et al., 2020; Hanson and Whiteheart, 2005; Ripstein et al., 2020; Rizo et al., 2019; Twomey et al., 2019; Wang et al., 2020). The aromatic residues of the six PL1 loops grip and translocate the peptide substrate in a sequence-promiscuous, hydrophobic milieu. Interestingly, *Nos*Rca and plant Rca do not have this canonical PL1 motif (Data S2). However, PL2 contains the highly conserved and functionally critical aromatic residue Tyr116 (Tyr188 in *Nt*Rca) (Stotz et al., 2011). In the ATP-bound subunits Rca2 to Rca6, Tyr116 hydrogen-bonds with Gln121 in the subsequent subunit, forming a network of interactions that rigidifies the central pore. In contrast, in the ADP-bound Rca1, PL2 is flexible and the position of Tyr116 is not well defined. Thus, it would appear that Tyr116 is unlikely to be involved in the process of peptide insertion in a manner similar to the Tyr in canonical pore-loops PL1. Instead, the formation of pockets between the staggered PL1 and PL2 of adjacent Rca subunits may be the critical feature for sequence specific substrate capture. As suggested for other AAA+ proteins, peptide threading may be mediated by sequential ATP hydrolysis around the hexamer ring (de la Pena et al., 2018; Dong et al., 2019; Fei et al., 2020; Ripstein et al., 2020; Rizo et al., 2019; Twomey et al., 2019; Wang et al., 2020).

### Function of SSUL Domain and AAA+ Core in Carboxysome Organization

Another important feature of *Nos*Rca is the presence of a Rubisco small subunit-like (SSUL) domain flexibly attached to the C-terminus of the AAA+ core in each Rca subunit. These SSUL domains mediate an alternative mode of interaction with Rubisco. They bind into a groove close to the equatorial region of Rubisco between the RbcL dimer units. This interaction has also been shown for the SSUL domains of the carboxysomal scaffolding protein CcmM (Wang et al., 2019), which mediates, via multivalent interactions, the formation of a Rubisco condensate for packaging into carboxysomes. *Nos*SSUL also contains the disulfide bond previously described for the SSUL module in CcmM, and thus its interaction with Rubisco is redox regulated, resulting in reduced affinity under oxidizing conditions.

In contrast to CcmM, which contains 3-5 SSUL modules as a linear fusion, *Nos*Rca contains six separate SSUL, one on each subunit, presumably providing high avidity for Rubisco. However, this arrangement may sterically preclude multivalent interactions required for efficient Rubisco network formation. Indeed, we found that critical valency is contributed by the AAA+ core of *Nos*Rca, as the mutational disruption of interface I resulted in a nearly complete loss in Rubisco condensate formation, despite the presence of the SSUL domains (Figure 6G). Thus, the seemingly unrelated interactions of *Nos*Rca in Rubisco repair and binding to Rubisco via the SSUL domains are linked and must cooperate to ensure Rca packing into carboxysomes (Figure 7E). Upon transfer into the oxidizing environment of the carboxysome, the SSUL-mediated association would then be released in favor of the functional interaction for Rubisco repair via the AAA+ core. It will be interesting to understand the interplay of *Nos*Rca and CcmM in Rubisco condensate formation during carboxysome biogenesis.

The acquisition of the SSUL domain by the *NosRca* AAA+ core probably occurred in the context of the appearance of carboxysomes about 350 million years ago, long after the primary endosymbiotic event leading to the evolution of chloroplasts (Rae et al., 2013). Efforts to introduce carboxysomes into chloroplasts in order to increase Rubisco efficiency (Hanson et al., 2016; Hennacy and Jonikas, 2020; Long et al., 2016; Rae et al., 2017), will have to consider coexpression of cyanobacterial Rca or attachment of the SSUL domain to plant activase.

## AUTHOR CONTRIBUTIONS

M.F. and H.W. planned and performed most of the experiments. M.F. established and performed all the activase assays and carried out turbidity assays together with H.W. M.F. was responsible for construct development and protein purification, and performed the synthesis of CABP. H.W. performed the cryo-EM structural analysis, including sample screening, grid preparation, data acquisition and single particle reconstructions, and fluorescence microscopy. L.P. contributed to cloning and protein purification, and obtained crystals of *Nos*RcaΔC and *Nos*SSUL. A.B. supervised the structural analyses and solved the crystal structures. M.H.-H. conceived the project and participated in data interpretation with F.U.H. and the other authors. M.H.-H and F.U.H wrote the manuscript with input from H.W., A.B. and M.F.

## ACKNOWLEDGMENTS

The technical assistance in the cloning and purification of proteins by S. Gaertner and R. Lange is gratefully acknowledged. We thank F. Bonneau and C. Basquin (Department of Structural Cell Biology) for help with the screening of a suitable buffer for protein purification, G. Thieulin-Pardo for providing *Nt*RcaR294V and *Nt*Rubisco as well as advice on Rca assays, R.H. Wilson for advice on expression of cyanobacterial Rubisco, X. Yan for advice on fluorescence microscopy, P. Wendler (University Potsdam) for initial analysis of *Nos*Rca by negative-stain EM, D. Cobessi (Institut de Biologie Structurale, Grenoble) for help with X-ray data collection of *Nos*SSUL and D. Bollschweiler for advice with cryo-EM data collection. We are grateful to the staff at FIP (French beamline for Investigation of Proteins) beamline BM30A at the European Synchrotron Radiation Facility (ESRF) and at beamline X06DA at the Swiss Synchrotron Light Source (SLS) in Villigen, Switzerland. Expert support by the Imaging, Crystallization and cryo-EM facilities of the Max Planck Institute of Biochemistry is acknowledged.

## METHODS

### Plasmids and Proteins

#### Plasmids

pET11a-*Nos*GroSEL was generated by amplifying the *groES-groEL* operon by PCR from genomic DNA of *Nostoc* sp. PCC 7120 (*Nos*GroSEL) and cloning into the pET11a vector via Gibson assembly. The Rubisco genes from *Nostoc* sp. PCC 7120 (*NosrbcL* and *NosrbcS*) were also amplified from genomic DNA by PCR and inserted into the pET28b vector. The His_6_-ubiquitin tag (H_6_ubi) was amplified from the pHUE plasmid (Baker et al., 2005; Catanzariti et al., 2004) and inserted as a N-terminal fusion to *NosrbcLS* to obtain pET28b-H_6_ubi-*Nos*LS. Finally, H_6_ubi-*Nos*LS was inserted C-terminally into pET11a-*Nos*GroSEL to obtain pET11a-*Nos*GroSEL-H_6_ubi-*Nos*LS. The N-terminal deletion mutant pET11a-*Nos*GroSEL-H_6_ubi-*Nos*LΔN12-S was generated by deleting the first 12 residues (MSYAQTKTQTKS) at the N-terminus of *Nos*RbcL by PCR. The plasmid pET11a-*At*Cpn60α/β-*At*Cpn20-*Nt*RbcL, a kind gift from the group of M.R. Hanson (Lin et al., 2019), was modified to generate pET11a-*At*Cpn60α/β-*At*Cpn20-H_6_ubi-*Nt*RbcLΔN9 by deletion of the first nine N-terminal residues (PQTETKASV) of *Nt*RbcL (*Nt*RbcLΔN9), and subsequently introducing the His_6_-ubiquitin tag (H_6_ubi) from the pHUE plasmid as a N-terminal fusion to *Nt*RbcLΔN9 (H_6_ubi-*Nt*RbcLΔN9).

pHUE-*Nos*Rca was generated by amplification of the *rca* gene from genomic DNA (Nostoc sp. PCC 7120) and subsequent cloning of residues 2-414 into the pHUE vector by using the SacII and EcoRI restriction sites. To generate pHUE-*Nos*RcaΔC, residues 292 and 293 in pHUE-*Nos*Rca were changed to two consecutive stop codons by site-directed mutagenesis. All mutations in pHUE-*Nos*Rca and pHUE-*Nos*RcaΔC were introduced by QuikChange mutagenesis (Agilent) resulting in the following constructs: pHUE-*Nos*Rca mutant (V91E/R92G/G93N/L244D/N248A/L250A); pHUE -*Nos*RcaΔC (V91E/R92G/G93N); pHUE - *Nos*RcaΔC (P140G/Y143A/D144A); pHUE -*Nos*RcaΔC (L244D/N248A/L250A).

pHUE-*Nos*Rca-SSUL was generated by cloning residues 325-414 (SSUL domain) of *Nos*Rca into the pHUE plasmid by PCR, and subsequent Gibson assembly (NEB).

#### Proteins

*Nt*Rubisco from *Nicotiana tabacum* leaves and *Nt*Rca(R294V) recombinantly expressed in *E. coli*, were purified as previously described (Servaites, 1985) (Stotz et al., 2011).

### Protein Expression and Purification

Protein concentrations were determined spectrophotometrically at 280 nm.

#### NosRcaΔC and NosSSUL for crystallography

*E. coli* BL21, harboring arabinose-inducible pBAD33-*Ec*GroSEL and pHUE-*Nos*Rca plasmids, was used for the expression and purification of *Nos*RcaΔC (residues 2-291) as a H_6_-ubiquitin (H_6_Ubi) fusion protein. Cells were grown in Luria-Bertani (LB) media at 37 °C/180 rpm until OD_600_ 0.3-0.4. The chaperonin GroES/EL was induced by addition of 1 % (w/v) L-arabinose. Two hours after induction, the temperature was reduced to 18 °C and expression of H_6_Ubi-NosRcaΔC was induced by addition of 0.5 mM isopropyl β-D-1-thiogalactopyranoside (IPTG) for 20 h. Cells were resuspended in ethanolamine (ETA) lysis buffer (50 mM ETA pH 8.0/300 mM NaCl/10 mM imidazole/5 % (v/v) glycerol) containing 1 mM PMSF, 10 mM 2-mercaptoethanol (2-ME), 1 g L^-1^ lysozyme and 5 U mL^-1^ benzonase, and lysed by sonication (15 x 15 s bursts with 75 s intermissions on ice). The supernatant obtained after high speed centrifugation (20 000 x g for 30 min at 4 °C) was loaded on a gravity nickel-nitrilotriacetic acid (Ni-NTA) metal affinity column (Qiagen), washed with 10 column volumes (CV) of ETA lysis buffer pH 9.2 containing 25 mM imidazole, and the protein eluted with ETA lysis buffer pH 9.2/200 mM imidazole. Fractions containing the protein were pooled and the H_6_Ubi moiety was cleaved by Usp2 (Baker et al., 2005; Catanzariti et al., 2004) at 4 °C overnight. After dialysis against 50 mM ETA pH 9.2/10 mM NaCl, *Nos*RcaΔC was loaded on a Mono Q HR 16/10 column (GE). The protein was eluted with a 10 CV gradient (0.01 – 0.5 M NaCl), concentrated and applied to a size-exclusion chromatography column (Superdex 200 10/300 GL; GE) equilibrated in 20 mM Tris-HCl pH 8.0/50 mM NaCl/5 mM MgCl_2_. Fractions containing the *Nos*RcaΔC were concentrated by ultrafiltration using Vivaspin MWCO 10000 (GE) and glycerol added to 5 % final prior to aliquoting and flash freezing in liquid N_2_. For all other studies the purification strategy was modified as stated below.

*Nos*SSUL (residues 325-414) was recombinantly expressed in *E. coli* from the pHUE-*Nos*Rca*-*SSUL plasmid as a H_6_Ubi fusion protein. Cells were grown in LB media at 30 °C/180 rpm until OD_600_ 0.3-0.4, and then *Nos*SSUL expression induced by addition of 0.2 mM IPTG and the cells shifted to 22 °C/120 rpm for 18 h. Cells were harvested and resuspended in ETA lysis buffer (50 mM ETA pH 9.2/300 mM NaCl/10 mM imidazole/5 % (v/v) glycerol) containing 1 mM PMSF, 10 mM 2-mercaptoethanol (2-ME), 1 g L^-1^ lysozyme and 5 U mL^-1^ benzonase, and lysed by sonication (15 x 15 s bursts with 75 s intermissions on ice). Purification of *Nos*SSUL was carried out essentially as above for *Nos*RcaΔC. Subsequent to dialysis after the first Ni-NTA column, *Nos*SSUL was applied to a second Ni-NTA resin to remove Usp2, H_6_Ubi and any uncleaved protein. The flow through was concentrated and applied to a size-exclusion chromatography column (HiLoad 16/60 Superdex 75; GE) equilibrated in 50 mM ETA pH 9.2/300 mM NaCl/1 mM DTT/5 % glycerol. Fractions containing the *Nos*SSUL were concentrated by ultrafiltration using Vivaspin MWCO 3000 (GE), aliquoted and flash frozen in liquid N_2_.

#### NosRca, NosRcaΔC and mutants

*Nos*Rca, *Nos*RcaΔC and mutants were expressed and purified from *E. coli* BL21 STAR cells harboring arabinose-inducible pBAD33-*Ec*GroSEL and the respective pHUE plasmid. Briefly, cells were grown in 2xYT media containing 10 mM KCl (Larimer and Soper, 1993) at 37 °C/180 rpm until OD_600_ 0.3-0.4. GroSEL was induced by addition of 0.4 % (w/v) L-arabinose. One hour after chaperonin induction, the temperature was reduced to 18 °C. When the cells had equilibrated to 18 °C (∼ 1 h), induction of protein was started by addition of 0.2 mM isopropyl β-D-1-thiogalactopyranoside (IPTG) and allowed to proceed for 18 h/120 rpm. Cells were harvested and incubated in buffer A (50 mM MMT pH 8.0/300 mM KCl/10 mM MgCl_2_/5 % glycerol) containing 1 g L^-1^ lysozyme/2.5 U mL^-1^/*Sm*DNAse//complete protease inhibitor cocktail (Roche) for 1 h prior to lysis using EmulsiFlex C5 (Avestin, Inc). Note MMT is a composite buffer consisting of DL-malic acid, MES monohydrate and Tris base in the molar ratio 1:2:2. After high speed centrifugation (40 000 x g/40 min/4 °C) the supernatant was loaded on to a gravity TALON metal affinity column (Takara), equilibrated and washed with 10 CV buffer A/20 mM imidazole. The bound protein was eluted with buffer A pH 8.4/200 mM imidazole, and diluted 3-fold in buffer A containing 5 mM 3-[(3-cholamidopropyl)dimethylammonio]-1-propanesulfonate (CHAPS)/10 % glycerol/5 mM 2-ME to a final protein concentration < 0.5 g L^-1^. The H_6_Ubi moiety was cleaved by Usp2 overnight at 10 °C. The cleaved protein was buffer exchanged on a HiPrep 26/10 desalting column (GE) to 50 mM MMT pH 8.4/10 mM KCl/5 % glycerol, and subsequently loaded on a MonoQ column (GE). The protein was eluted with a 10 CV gradient (0.01 – 0.5 M KCl), concentrated to ∼5 mL and applied onto a size-exclusion chromatography column (HiLoad 16/60 Superdex 200; GE) equilibrated in buffer B (50 mM MMT pH 8.4/100 mM KCl/10 mM MgCl_2_/5 % glycerol). The protein containing fractions were concentrated by ultrafiltration using Vivaspin MWCO 3000 (GE), aliquoted and flash frozen in liquid N_2_. To generate reduced *Nos*Rca, 5 mM DTT was added to all buffers after Usp2 cleavage. The oxidized *Nos*Rca purified in the absence of DTT was allowed to further air oxidize on ice for 8 h after the final column prior to concentrating, aliquoting and flash freezing.

#### NosRubisco and mutants

*Nos*Rubisco was expressed in *E. coli* BL21 STAR cells harboring the IPTG-inducible *Nostoc* chaperonin on pET11a-*Nos*GroSEL-H_6_ubi-*Nos*LS plasmid, in which the cleavable H_6_Ubi motif is attached at the N-terminus of *Nos*RbcL. Cells were grown in 2 x YT/10 mM KCl media (Larimer and Soper, 1993) at 37 °C/180 rpm until OD_600_ 0.6 – 0.8. *Nos*GroES/EL and H_6_ubi-*Nos*RbcLS were induced by addition of 0.5 mM IPTG and cells shifted to 22 °C at 120 rpm for 22 h. Cells were lysed, cell debris removed and the supernatant loaded onto a TALON resin column as described above. After 10 CV washes in buffer A/20 mM imidazole the H_6_Ubi-tagged Rubisco complex was eluted with buffer A pH 8.4/200 mM imidazole. Usp2-mediated digestion was performed overnight in presence of 5 mM 2-ME at 10 °C. After removal of imidazole on a HiPrep 26/10 desalting column (GE) equilibrated in buffer A, the buffer-exchanged protein eluate was applied to a TALON resin column for removal of Usp2, the cleaved H_6_Ubi moiety and any uncleaved protein. The flow through was concentrated to ∼ 5 mL and applied onto a size-exclusion chromatography column as above, equilibrated in buffer B containing 30 mM NaHCO_3_. The *Nos*Rubisco containing fractions were concentrated by ultrafiltration using Vivaspin MWCO 30000 (GE), aliquoted and flash frozen in liquid N_2_.

#### NtRubiscoΔN

*Nt*RubiscoΔN, with 9 residues deleted at the N-terminus of the RbcL subunit, was expressed in *E. coli* BL21 STAR cells harboring the IPTG-inducible plasmids pET11a-*At*Cpn60αβ-*At*Cpn20-H_6_ubi-ΔN9*Nt*RbcL and pCDF-*Nt*XSR_1_R_2_B_2_ (kind gift from MR Hanson). Cells were grown in 0.1 L *ZYP-5052* auto-induction media (Lin et al., 2019; Studier, 2005) at 37 °C/160 rpm for 6 h. IPTG induction was performed for at 23 °C/120 rpm for 22 h. Harvested cells were re-suspended and incubated in buffer A containing 1 g L^-1^ lysozyme, 2.5 U mL^-1^ *Sm*DNAse, complete protease inhibitor cocktail (Roche) for 1 h prior to lysis by sonication. After high speed centrifugation (40 000 x g/40 min/4 °C) the supernatant was loaded on a HiTrap (GE) TALON Crude (Takara) metal affinity column, washed with 10 CV of buffer A prior to elution by a 10 CV linear imidazole gradient 0 to 200 mM for separation of H_6_Ubi-tagged Rubisco from *At*Cpn60αβ. Fractions containing only H_6_Ubi-tagged ΔN9*Nt*Rubisco were selected by immunoblotting against *Nt*RbcL and *At*Cpn60α/β. The pooled fractions were subjected to digestion by Usp2 (8 h/10 °C) in the presence of 5mM 2-ME. The reaction was then buffer exchanged to 50 mM MMT pH 8.4/100 mM KCl/5 % glycerol on a HiPrep 26/10 desalting column (GE), and applied to a gravity TALON (Takara) metal affinity column for removal of Usp2, the cleaved H_6_Ubi moiety and any uncleaved protein. The flow through was concentrated by ultrafiltration using Vivaspin MWCO 30000 (GE), aliquoted and flash frozen in liquid N_2_.

### CABP Synthesis

2-Carboxyarabinitol-1,5-diposphate (CABP) was synthesized according to (Pierce et al., 1980). In brief, 100 μmol RuBP was incubated with a 2-fold molar excess of KCN in 5 mL 0.1 M Tris-acetate pH 8.3 for 48 h at 25 °C. The racemic mixture of 2-carboxyribitol-1,5-diphosphate (CRBP) and CABP was treated with the cation exchange resin AG50W-X8 (H^+^), filtered and freeze dried. To separate the enantiomers, the lactonized products were dissolved in 3 mM HCl and applied to a 120 mL AG1X8 (Cl^-^) column equilibrated in 3 mM HCL, eluted over a 4 L gradient from 0 – 0.4 M LiCl, and then the 50 mL fractions were assayed for total phosphate (Chifflet et al., 1988). CABP containing fractions were pooled and reduced to a volume of 50 mL in a rotary evaporator at 30 °C. Addition of 3-fold molar excess of barium acetate precipitated the CABP as barium salt (1 h at −20 °C). The precipitate was collected by centrifugation (5000 x g for 20 min at 4 °C) and redissolved by addition of acid-washed AG50W-X8 resin. After filtration, the purified CABP was freeze-dried and redissolved in 50 mM Bicine-NaOH pH 9.3. Complete saponification was ensured by incubation on ice for 24 h, before aliquoting and flash freezing in liquid N_2_.

### Enzymatic Assays

All assays were performed at 25 °C

#### Rubisco reactivation

CO_2_ fixation by Rubisco and Rubisco activase activity were measured as previously described (Barta et al., 2011; Esau et al., 1996) in buffer 50 mM MMT pH 8.4/50 mM KCl/30 mM NaH^14/12^CO_3_ (14 Bq nmol^-1^)/10 mM MgCl_2_/3 mM phosphocreatine/50 U mL^-1^ phosphocreatine kinase containing RuBP and ATP as indicated in the figure legends. The carbamylated Rubisco (ECM) was obtained by preincubating Rubisco in 100 mM MMT pH 8.4/60 mM NaHCO_3_/20 mM MgCl_2_ for 10 min. The uncarbamylated Rubisco (E) was obtained by buffer exchange of the purified Rubisco into 100 mM MMT pH 8.4/50 mM KCl/4 mM EDTA. The inhibited *Nos*Rubisco (E.XuBP, E.RuBP and ECM.CABP) and inhibited *Nt*Rubisco (E.RuBP) were formed by addition of the respective inhibitory sugars and incubation for 30-60 min. Time course experiments were initiated by addition of ECM, ECM.CABP, E, E.RuBP or E.XuBP (0.25 μM active sites) to the reaction assay including RuBP (0.4 mM unless otherwise indicated), Rca and ATP (3 mM) as indicated.

Dose dependent reactivation of *Nos*Rubisco (ECM.CABP) was measured in presence of ATP (3 mM) and at the indicated *Nos*RcaΔC concentrations (0.125, 0.25, 0.5, 1.25 μM hexamer). The single time-point for the amount of CO_2_ fixed was analyzed at 8 min after initiation of the reaction and conducted in the presence of 1 mM RuBP to ensure steady state kinetics for the ECM control.

To estimate specific activity and account for background counts, corresponding samples were measured in 5 or 6 replicates per assay.

#### ATPase assay

ATP hydrolysis was enzymatically coupled to NADH oxidation and measured spectrophotometrically at 25 °C (Barta et al., 2011; Mueller-Cajar et al., 2011). The assay was started by addition of *Nos*Rca, *Nos*RcaΔC, or mutants (*Nos*Rca V91E/R92G/G93N/ L244D/N248A/L250A, *Nos*RcaΔC P140G/Y143A/D144A, *Nos*RcaΔC V91E/R92G/G93N, *Nos*RcaΔC L244D/N248A/L250A) at 0.5 μM (hexamer) in assay buffer (100 mM MMT pH 8.4/20 mM KCl/10 mM MgCl_2_/5 mM DTT/2 mM phosphoenolpyruvate/0.3 mM NADH/2 mM ATP/∼ 30 U mL^-1^ pyruvate kinase/45 U mL^-1^ lactic dehydrogenase). ECM and ECM.CABP (0.25 μM hexadecamer) were prepared as described above and added as indicated. To suppress turbidity due to condensate formation of activated or inhibited Rubisco (ECM or ECM.CABP, respectively) in the presence of *Nos*Rca, the salt concentration was increased to 200 mM KCl in the assay buffer.

### Turbidity Assay

Measurements were performed at 25 °C in buffer C (50 mM MMT pH 8.4/50 mM KCl/10 mM MgCl_2_) for oxidized *Nos*Rca and in the presence of additional 5 mM DTT for reduced *Nos*Rca /*Nos*RcaΔC /*Nos*Rca-IF_mut_. Reactions (100 μL) containing Rubisco (0.25 μM) and different concentrations of *Nos*Rca or *Nos*RcaΔC (0.5 μM) or *Nos*Rca-IF_mut_ (0.5 μM) in the absence or presence of 2 mM ATPγS as indicated in the figure legends were mixed rapidly by vortexing, and absorbance at 340 nm was monitored as a function of time on a Jasco V-560 spectrophotometer.

### Liquid-liquid Phase Separation

For analysis of LLPS, Rubisco holoenzyme was labeled at the N terminus with the fluorophore Alexa Fluor 532 NHS ester (ThermoFisher) according to manufacturer’s instructions (∼2 dye molecules bound per Rubisco holoenzyme). Reduced *Nos*Rca, *Nos*RcaΔC and *Nos*Rca-IF_mut_ was labelled at the N terminus with the fluorophore Alexa Fluor 405 NHS ester (ThermoFisher) (∼2.2, 1.7 and 3 dye molecules bound per Rca hexamer, respectively). Labeled protein was mixed with unlabeled protein, at a ratio of 1:10. Reactions (20 μL) in buffer C containing 5 mM DTT with reduced *Nos*Rca, *Nos*RcaΔC or *Nos*Rca-IF_mut_ (0.5 μM) and Rubisco (0.25 μM) were incubated for 5 min at 25 °C and then transferred to an uncoated chambered coverslip (μ-Slide angiogenesis; Ibidi) for another 5 min before analysis. Images were illuminated with a Lumencor SPECTRA X Light Engine at 398 nm and 558 nm for fluorescence imaging. Images were recorded by focusing on the bottom of the plate using a Leica Thunder Widefield 2 microscope with Leica DFC9000 GTC camera and a HC PL APO 63x/1.47 oil objective.

### Size-exclusion Chromatography Coupled to Multi-angle Static Light Scattering (SEC-MALS)

Purified proteins at 2 mg mL^-1^ was analyzed using static and dynamic light scattering by auto-injection of the sample onto a SEC column (5 μm, 4.6×x300 mm column, Wyatt Technology, product # WTC-030N5) at a flow rate of 0.2 mL min^-1^ in buffer 50 mM MMT pH 8.4/100 mM KCl/10 mM MgCl_2_ at 25 °C in the presence or absence of nucleotide (1 mM). The column was in line with the following detectors: a variable UV absorbance detector set at 280 nm (Agilent 1100 series), the DAWN EOS MALS detector (Wyatt Technology, 690 nm laser) and the Optilab rEX^TM^ refractive index detector (Wyatt Technology, 690 nm laser) (Wyatt, 1993). Molecular masses were calculated using the ASTRA software (Wyatt Technology) with the dn/dc value set to 0.185 mL g^-1^. Bovine serum albumin (Thermo) was used as the calibration standard.

### Electron Microscopy and Reconstruction

#### Cryo-EM

All cryo-grids were prepared with a Vitrobot Mark 4 (FEI). A volume of 3 μL of the sample was applied to a glow-discharged grid (Quantifoil R2/1 300 mesh) at 25 °C and 90 % humidity, then semi-automatically blotted and plunge-frozen into liquid ethane.

To capture the interaction between *Nos*RcaΔC and CABP-inhibited Rubisco (*Nos*RcaΔC:Rubisco), *Nos*Rubisco (1 μM) was first carbamylated by incubation in buffer C containing 10 mM NaHCO_3_ for 10 min at 25 °C, then inhibited with 8 μM CABP at 25 °C for 1 h. The *Nos*RcaΔC:Rubisco (*Nos*RcaΔC:ECM.CABP) complex was formed following a published nucleotide-substitution strategy (Dong et al., 2019). Specifically, ECM.CABP (0.5 μM) was incubated with *Nos*RcaΔC (10 μM) at 25 °C in the presence of ATP (2 mM) for 10 s, followed by the addition ATPγS (2 mM), and incubated at 25 °C for another 10 min before preparing the cryo-grids as stated above. The cryo-grids were initially screened on a Talos Arctica (FEI) transmission electron microscope (TEM). Selected grids were transferred to a Titan Krios 300 kV TEM (FEI) equipped with GIF Quantum Energy Filters (Gatan), and a K3 direct detector (Gatan). 9,042 movies were automatically collected by SerialEM (Mastronarde, 2005) using a pixel size of 0.8512 Å. The total exposure time of 2.8 s was divided into 31 frames with an accumulated dose of 60 electrons per Å^2^ and a defocus range of −0.7 μm to −2.5 μm.

For Preparing the *Nos*Rca:Rubisco complex, *Nos*Rubisco (1.25 μM) was mixed with *Nos*Rca (15 μM) in buffer C containing 5 mM DTT and cryo-grids prepared as above. The cryo-grids were screened on a Glacios transmission electron microscope (Thermo Scientific), equipped with K2 summit direct electron detector (Gatan), operated at 200 keV. Selected grid on stage was used for data collection directly with K2 summit. Exposure times of 12 s were divided into 40 frames with an accumulated dose of 47 electrons per Å^2^. 1,570 movies were automatically collected by SerialEM (Mastronarde, 2005) with a pixel size of 1.885 Å and a defocus range of −1 μm to −4.5 μm.

#### Image processing

For the *Nos*RcaΔC:Rubisco (*Nos*RcaΔC:ECM·CABP) dataset, on-the-fly processing during data collection was performed with MotionCorr2 and CTFFIND-4.1, as implemented in the Focus software (Biyani et al., 2017). Only micrographs with good particle quality, with an estimated maximum resolution below 5 Å, were kept for further data processing with RELION 3.0. A total of 519,151 particles were auto-picked by Gautomatch (http://www.mrc-lmb.cam.ac.uk/kzhang/Gautomatch) and extracted at a pixel size of 3.4048 Å (four-fold binned). The first round of 2D-classification was used to exclude ice contaminations and classes with no structural features. The selected particles were next refined according to a Rubisco reference converted from the dataset of PDB 1RBL (Newman et al., 1993), re-centered, and then subjected to a second round of 2D-classification. Free *Nos*RcaΔC classes were excluded, which resulted in the selection of 106,831 particles (Figure S3). 3D-classification with the Rubisco reference showed only one class of the *Nos*RcaΔC hexamer in complex with Rubisco, containing 30,607 particles. These particles were re-extracted at full resolution (0.8512 Å pixel size) and subjected to “polishing” with RELION to generate “shiny” particles. To deal with multiple *Nos*RcaΔC hexamers bound per Rubisco, we followed a previously published symmetry-expansion procedure (Wang et al., 2019). In detail, particles were first aligned with *D4* symmetry. Then *D4* symmetry was released and yielded 8-fold particles by the symmetry-expanding command, relion_particle_symmetry_expand. A focused 3D-classification in *Nos*RcaΔC with symmetry-released particles revealed one class with *Nos*RcaΔC hexamer occupancy, comprising of 21,149 particles (Figure S3). These particles generated the final map of the *Nos*RcaΔC:ECM·CABP complex at 2.86 Å resolution, determined by gold-standard Fourier shell correlation (FSC) with a cutoff at 0.143. Particle subtraction of the Rubisco signal improved the alignment accuracy of the *Nos*RcaΔC hexamer, and generated a local *Nos*RcaΔC map at 3.29 Å resolution.

The raw movies of the *Nos*Rca:Rubisco dataset were first processed with MotionCorr2 (Zheng et al., 2017) with dose-weighting. CTFFIND-4.1(Rohou and Grigorieff, 2015) estimated the CTF parameters for each micrograph. 298,336 Particles were picked by Gautomatch (http://www.mrc-lmb.cam.ac.uk/kzhang/Gautomatch). Two rounds of 2D-classification excluded ice contaminations and classes with no structural features, and resulted in 45,859 clean particles (Figure S7). These particles were used to generate a reference for 3D-classification in RELION 3.0. (Scheres, 2012) with the 3D initial model module. 3D-classification revealed one class with detailed Rubisco features (27,527 particles). To deal with multiple SSUL domains bound per Rubisco, we followed the same symmetry-expansion procedure previously reported (Wang et al., 2019). Particles were first aligned with *D4* symmetry. Each asymmetric unit L_2_S_2_SSUL was processed as an individual particle, which is achieved by the symmetry-expanding command, relion_particle_symmetry_expand, and particle subtraction. A focused classification with a SSUL mask resulted in one class of particles with detailed SSUL feature. 32,128 particles from this class were selected and subjected to final local refinement. Post-processing improved the map resolution to 8.2 Å.

#### Model building

*Nos*RcaΔC:ECM·CABP – *Nos*RcaΔC model building was initiated by rigid-body fitting the *Nos*RcaΔC subdomains from the crystal structure into the cryo-EM density, followed by manual editing using Coot (Emsley and Cowtan, 2004). This model was refined in reciprocal space with REFMAC5 (Murshudov et al., 2011). The ECM.CABP model was generated with SWISS-MODEL (Waterhouse et al., 2018) based on the coordinates of Rubisco from *Chlamydomonas reinhardtii* (PDB 1UZH) (Karkehabadi et al., 2005). This model was placed into the cryo-EM density using Chimera (Pettersen et al., 2004), followed by manual editing using Coot. Residues with disordered side-chains were truncated at C-β. This model was refined in reciprocal space with REFMAC5 (Murshudov et al., 2011), using non-crystallographic symmetry restraints.

*Nos*Rca:Rubisco – First, the crystal structures of thiol-reduced *Nos*SSUL and *Nos*Rubisco were placed into the density using Chimera, followed by manual editing using Coot. The resulting model was refined in reciprocal space with REFMAC5, using jelly-body restraints. The used structure factors were calculated from a masked map.

The models and the electron density maps for *Nos*RcaΔC:Rubisco and *Nos*Rca Rubisco have been deposited to the wwPDB database under PDB/EMDB accession codes 6Z1F/EMD-11028 and 6Z1F/EMD-11029, respectively.

### Crystallization and Data Collection

The *Nos*RcaΔC construct used for crystallization includes residues 2–291. The *Nos*SSUL construct used for crystallization includes residues 325–414.

*Nos*RcaΔC – Crystals were grown by the hanging-drop vapor diffusion method at 4 °C. Drops containing 2 μL of a 1:1 mixture of 4.9 mg mL^-1^ *Nos*RcaΔC in buffer 20 mM Tris-HCl pH 8.0/50 mM NaCl/5 mM MgCl_2_ and precipitant were equilibrated against 500 μL precipitant. The precipitant contained 2.2 or 2.3 M Na-acetate pH 7.0.

*Nos*SSUL – Crystals were grown by the hanging-drop vapor diffusion method at 4 °C. Drops containing 3 μL of a 1:1 mixture of 5.2 mg mL^-1^ *Nos*SSUL in buffer 50 mM ETA pH 9.2/300 mM NaCl/1 mM DTT/5 % glycerol and precipitant were equilibrated against 500 μL precipitant. The precipitant contained 26 % PEG-3350 and 50 mM MES-NaOH pH 6.0.

For cryo-mounting, the crystals were transferred into a cryo-solution that was precipitant containing 15 % glycerol in addition and subsequently cryo-cooled by dipping into liquid nitrogen.

#### Crystallographic data collection, structure solution and refinement

*Nos*RcaΔC – The diffraction data of the crystals of *Nos*RcaΔC were collected by the oscillation method at beamline X06DA at the Swiss Synchrotron Light Source (SLS) in Villigen, Switzerland. The diffraction data from *Nos*RcaΔC crystals were integrated with XDS and further processed with POINTLESS (Evans, 2006), SCALA (Evans, 1997) and CTRUNCATE (French and Wilson, 1978). The structure was solved at 3.4 Å from a Gadolinium (GdCl_3_) derivative by Gd-multi-wavelength anomalous diffraction (MAD) using ShelxC/D/E (Sheldrick, 2010) as implemented in the Hkl2map GUI (Pape and Schneider, 2004). The seven Gd sites were refined and phases calculated with SHARP (de la Fortelle and Bricogne, 1997). The map, calculated after density modification with RESOLVE (Terwilliger, 2000), assuming a solvent content of 55 %, revealed features of secondary structure elements. A preliminary model was auto-built with Buccaneer (Cowtan, 2006), and missing portions added manually using COOT (Emsley and Cowtan, 2004). REFMAC5 was used for initial model refinement (Murshudov et al., 2011). The final refinement was performed with phenix.refine (Adams et al., 2010) using translation-libration-screw (TLS) parametrization of B-factors. The native structure was solved by molecular replacement. The final models contain two copies of *Nos*RcaΔC per asymmetric unit. One chain (chain B) has ADP bound. Residues 276–291 were disordered in chain A. In chain B, residues 105–115, 248–254 and 279–291 are missing. Residues facing solvent channels with disordered side-chains were modelled as alanine. The model of the native structure contains 14 ordered water molecules and exhibits reasonable stereochemistry with 96.6 % of the residues in the favored regions of the Ramachandran plot according to the criteria of MolProbity (Chen et al., 2010).

*Nos*SSUL – The native diffraction data of the crystals of *Nos*SSUL were collected at the automated beamline ID30A-1 at the European Synchrotron Radiation Facility (ESRF) in Grenoble, France. MAD data for a presumed Pt-derivative were collected at beamline BM30A at ESRF. The diffraction data from *Nos*SSUL crystals were integrated with XDS and further processed with POINTLESS (Evans, 2006), AIMLESS (Evans and Murshudov, 2013) and CTRUNCATE (French and Wilson, 1978) as implemented in the CCP4i graphical user interface (Potterton et al., 2003). The structure of *Nos*SSUL was solved by MAD using the Auto-Rickshaw platform (Panjikar et al., 2005). The anomalous scatterers were a bound Ni^2+^ atom from protein purification and presumably ordered Cl^−^ ions. The chemical environment of the sites was not compatible with PtCl_4_^2−^. The asymmetric unit contained two copies of the SSUL domain. The model was edited manually using Coot (Emsley and Cowtan, 2004). REFMAC5 was used for model refinement (Murshudov et al., 2011). The model contains 203 ordered water molecules and exhibits reasonable stereochemistry with 99.5 % of the residues in the favored regions of the Ramachandran plot according to the criteria of MolProbity (Chen et al., 2010).

Figures were created with PyMol (http://www.pymol.org/) and ESPript (Gouet et al., 1999).

### Structure Analysis

The quality of the structural models was analyzed with the program Molprobity (Chen et al., 2010). Coordinates were aligned with Lsqkab and Lsqman (Kleywegt and Jones, 1994). Molecular interfaces were analyzed with PISA (Krissinel and Henrick, 2007) and Contact, as implemented in the CCP4i graphical user interface (Potterton et al., 2003). Figures were created with Chimera (Pettersen et al., 2004), PyMol (http://www.pymol.org/) and ESPript (Gouet et al., 1999).

### Data Resources

The models and the electron density maps for *Nos*RcaΔC:Rubisco and *Nos*Rca Rubisco have been deposited to the wwPDB database under PDB/EMDB accession codes 6Z1F/EMD-11028 and 6Z1F/EMD-11029, respectively. The crystallographic models and structure factors for *Nos*RcaΔC-Gd complex, *Nos*RcaΔC and *Nos*SSUL have been deposited to the PDB database under accession codes 6Z1D, 6Z1E and 6HAS, respectively.

## SUPPLEMENTAL FIGURES

**Figure S1.**
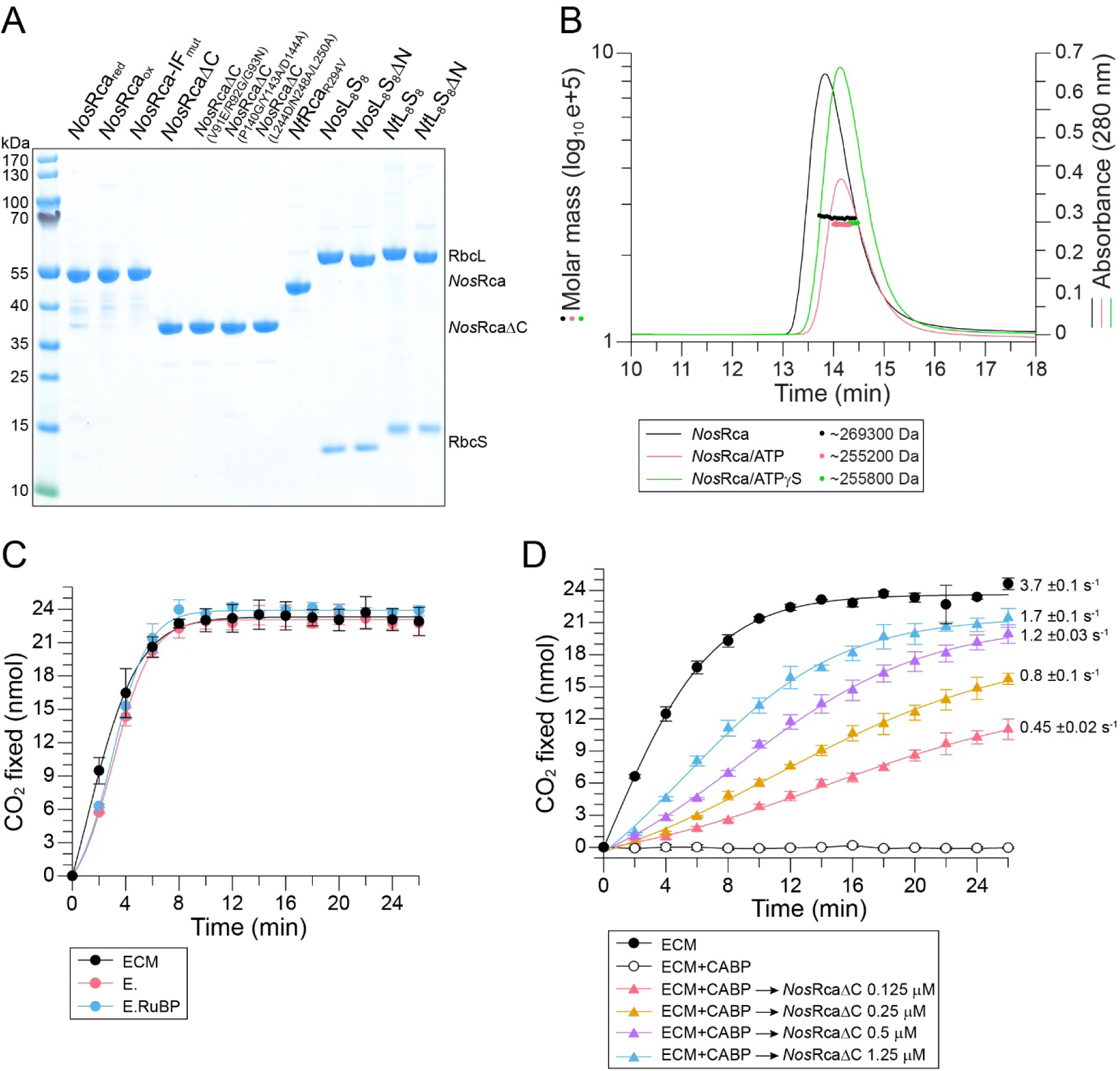
Rubisco Activase Function of *Nos*Rca. (A) Purified proteins used in this study. Protein concentrations were determined spectrophotometrically. 1.5 μg protein was analyzed by SDS-PAGE. (B) *Nos*Rca is a hexamer in solution. SEC-MALS analysis of *Nos*Rca in the absence or presence of nucleotide (1 mM ATP or ATPγS). The chromatographic absorbance traces at 280 nm wavelength are shown. The molecular mass determined for the protein peaks by static light scattering is indicated. (C) Non-cambamylated *Nos*Rubisco is not inhibited by RuBP. CO_2_ fixation assays were performed with carbamylated (ECM), non-carbamylated (E) and non-carbamylated Rubisco with RuBP (E.RuBP) as in Figure 1C. Error bars represent SD of at least three independent replicates. (D) Dependence of Rubisco reactivation on *Nos*RcaΔC concentration. CABP inhibited *Nos*Rubisco (ECM.CABP) was incubated with increasing concentrations of *Nos*RcaΔC (0.125, 0.25, 0.5 and 1.25 μM hexamer) in the presence of 3 mM ATP, and CO_2_ fixation measured as in Figure 1D. Approximate rates of CO_2_ fixation were determined from the linear parts of the curves. Error bars represent SD of three independent replicates.

**Figure S2.**
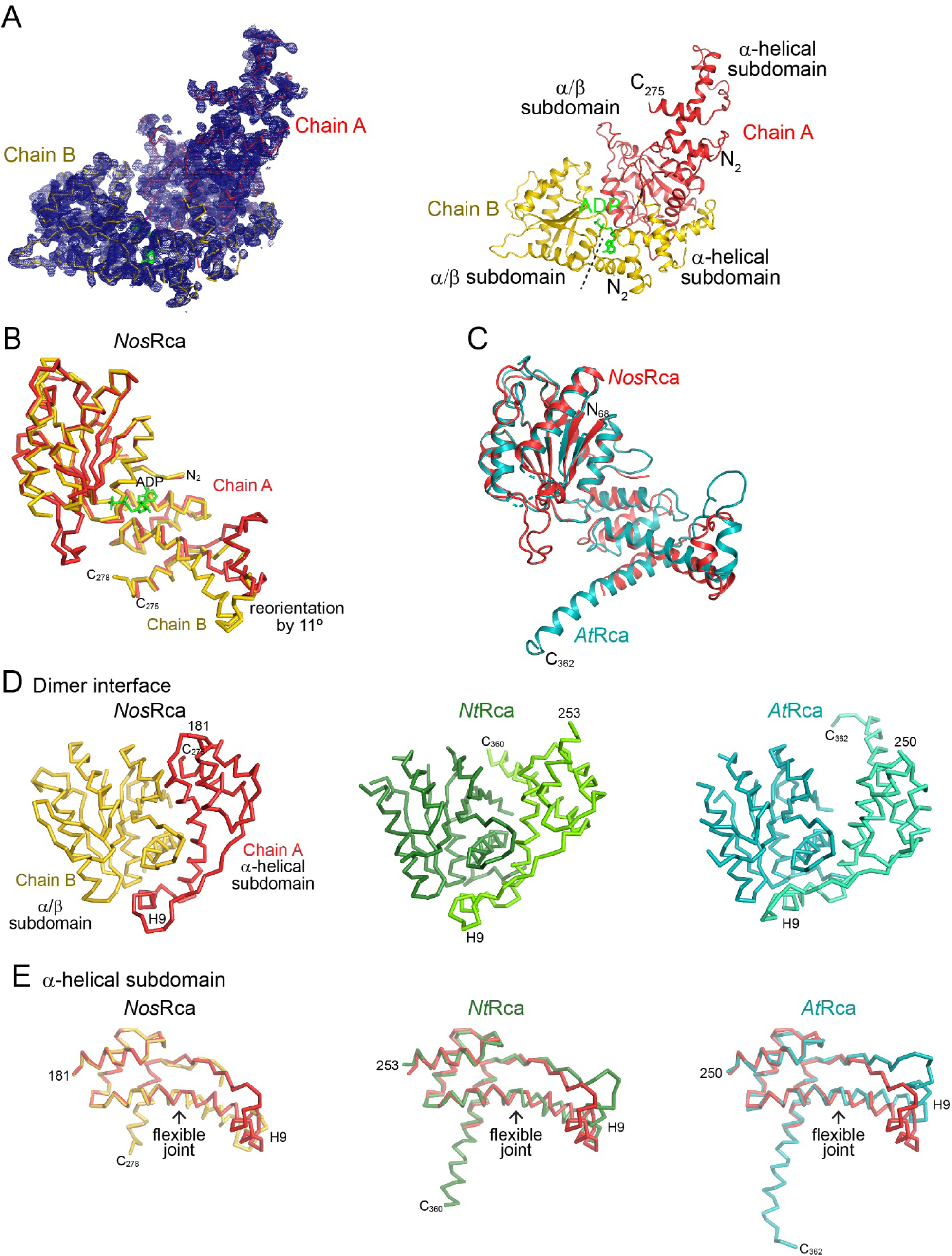
Crystal Structure of *Nos*RcaΔC. (A) Left: Unbiased experimental electron density at 2.7 Å resolution for the Gd^3+^ complex of *Nos*RcaΔC (for data collection and refinement statistics, see Table S2). The density after Gd-MAD phasing and density modification contoured at 1.5 σ is shown as a blue meshwork. Chains A and B of the asymmetric unit are shown as Cα traces in red and yellow, respectively. ADP is represented as a wire-frame model in green. Right: Final model of the asymmetric unit in the *Nos*RcaΔC crystal lattice. The two crystallographically independent chains of *Nos*RcaΔC are shown as ribbons. Subdomains and chain termini are indicated. The dashed line indicates the subdomain boundary in chain B. (B) Superposition of chain A with chain B in the asymmetric unit of *Nos*RcaΔC. Protein chains are represented as Cα traces. (C) Superposition of chain A (no nucleotide bound) of the asymmetric unit of *Nos*RcaΔC with *At*Rca (PDB: 4W5W) (Hasse et al., 2015). Chain A of *Nos*RcaΔC and *At*Rca are shown as ribbons in red and teal, respectively. Chain termini are indicated. (D) A conserved subunit-subunit interaction found in *Nos*Rca (left), *Nt*Rca (middle) and *At*Rca (right). The interaction is formed between the α/β subdomain in one subunit and the C-terminal half of helix α8, α9 (H9) and the connecting linker in the α-helical subdomain of the adjacent subunit. Adjacent subunits in the *Nt*Rca crystal structure are shown in dark and bright green, and in teal and cyan for *At*Rca. The location of helix H9 and chain termini are indicated. Note that the orientation of helix α9 (H9) with respect to the four-helix bundle in the α-helical subdomain differs between the structures; the long helix α8 and the long α9-α10 linker act as a stalk that can twist and bend like a flexible joint. (E) Superposition of the α-helical subdomain of chain A in *Nos*Rca with chain B (left), with *Nt*Rca (middle) and *At*Rca (right). The r.m.s.d. values for matching Cα positions are 0.24 Å (57 Cα positions), 0.58 Å (42 Cα positions) and 0.55 Å (47 Cα positions), respectively. The locations of helix α9 (H9) and the flexible joint region are indicated. Chain termini are indicated.

**Figure S3.**
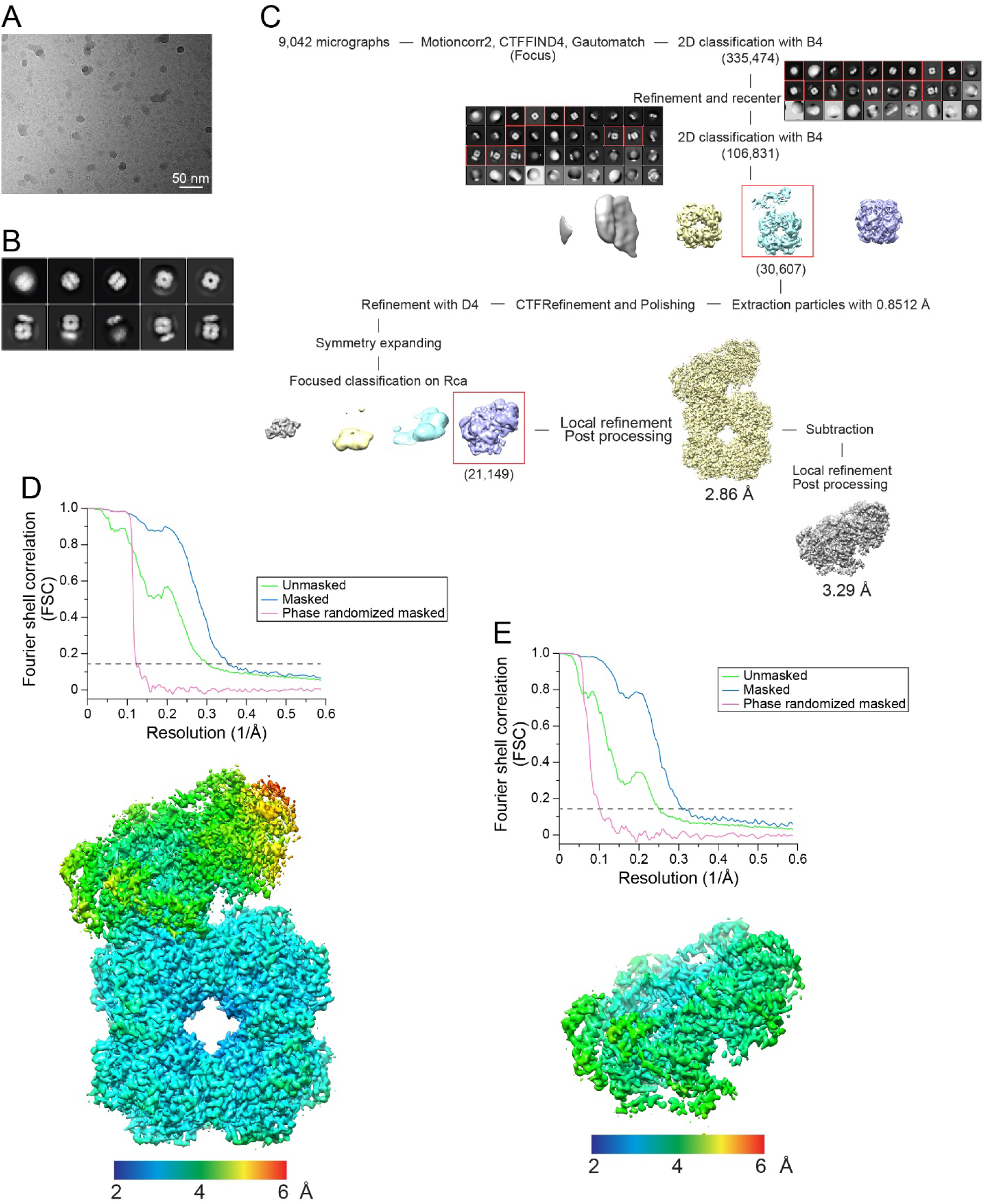
Cryo-EM Single-particle Reconstruction of *Nos*RcaΔC:Rubisco Complex. (A) A representative micrograph of *Nos*RcaΔC:Rubisco complexes. (B) 2D class averages of complexes in (A). (C) The single-particle data processing workflow for the *Nos*RcaΔC:Rubisco complex. Particle numbers are in parentheses. B4, 4×4 pixel-binned image. See STAR Methods for details. (D and E) Gold-standard FSC curves and local resolution maps of the *Nos*RcaΔC:Rubisco reconstruction (D) and the *Nos*RcaΔC local map (E). The resolutions are ∼2.86 Å and ∼3.29 Å, respectively, at the FSC cutoff of 0.143 for the masked and B-factor sharpened curves. The color gradient from blue to red indicates local resolution from 2.0 to 6.0 Å.

**Figure S4.**
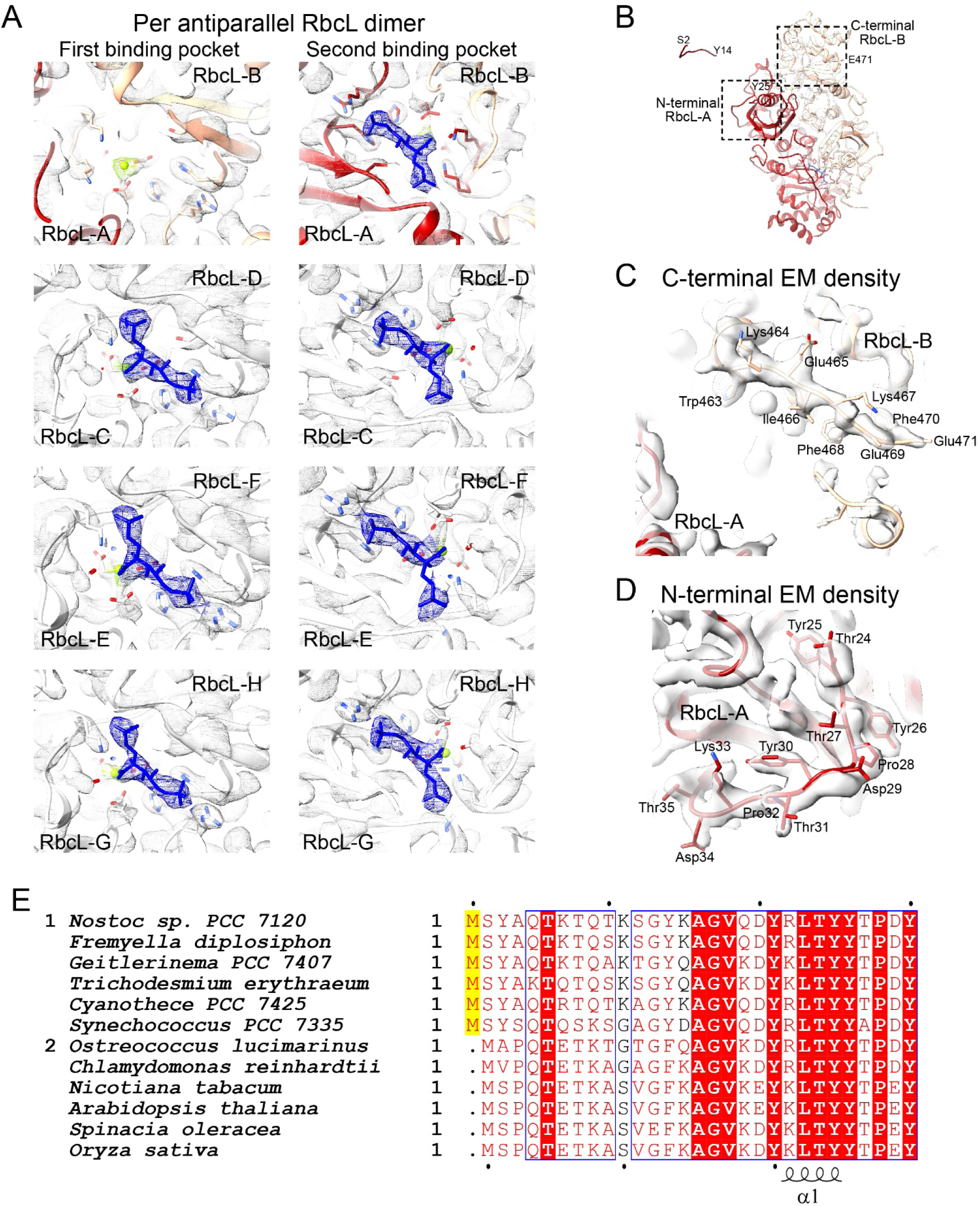
Key Structural Features of Inhibited *Nos*Rubisco in Complex With *Nos*RcaΔC. (A) Close-up of the cryo-EM density of the eight substrate binding pockets of *Nos*Rubisco. Each RbcL anti-parallel dimer has two binding pockets. The density is represented as a transparent iso-contour surface. CABP density is shown as a meshwork in blue. Protein is represented by ribbons with ligand-interacting sidechains and CABP in stick representation. Mg^2+^ ions are shown as green spheres. (B) An overview of the RbcL-A/RbcL-B dimer that is engaged by *Nos*RcaΔC. RbcL-A and RbcL-B are shown in red and peach, respectively. The N- and C-terminal residues, as well as the RbcL-A N-terminal peptide bound in the hexamer pore of *Nos*RcaΔC are indicated. (C and D) Zoomed-in views of the N- and C-terminal regions, indicated by dotted boxes in (B). The EM densities of the C-terminal residues of RbcL-B (C) and the N-terminal residues of RbcL-A (D) are well-resolved in the cryo-EM density map of the *Nos*RcaΔC:Rubisco complex. (E) Alignment of the N-terminal sequences of RbcL from selected cyanobacterial and eukaryotic form 1B Rubisco proteins. Similar residues are shown in red and identical residues in white on a red background. Blue frames indicate homologous regions. Yellow background indicates systematic differences between cyanobacterial and eukaryotic sequences. The secondary structure elements of RbcL from *Oryza sativa* (PDB: 1WDD) (Matsumura et al., 2012) are indicated below the sequences. The Uniprot and Genebank accession codes for the sequences are: P00879, *Nostoc* PCC 7120; NZ JH930358.1, *Fremyella diplosiphon*; K9SCF1, *Geitlerinema* PCC 7407; Q10WH6, *Trichodesmium erythraeum*; B8HQS5, *Cyanothece* PCC 7425; B4WP00, *Synechococcus* PCC 7335; Q0P3J3, *Ostreococcus lucimarinus*; P00877, *Chlamydomonas reinhardtii*; P00876, *Nicotiana tabacum*; O03042, *Arabidopsis thaliana*; P00875, *Spinacia oleracea*; P0C510, *Oryza sativa*.

**Figure S5.**
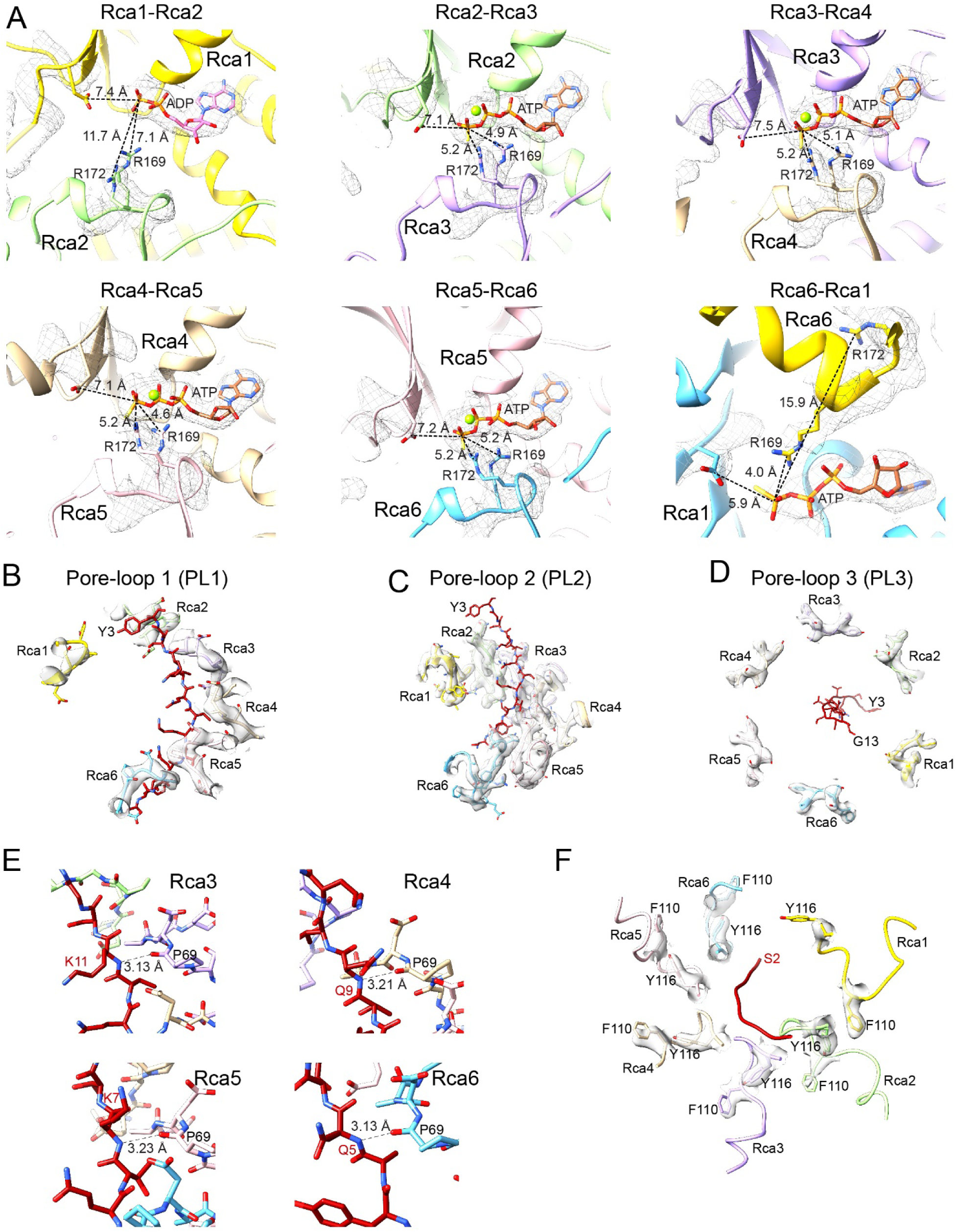
Key Structural Features in *Nos*RcaΔC in the Complex with *Nos*Rubisco. (A) Zoomed-in views of the nucleotide binding pockets formed between adjacent *Nos*Rca subunits. Distances of the carbon atom in the guanidino group of Arg169 and Arg172, and of the carboxylate carbon of Asp102 (Walker B) to the last phosphorus atom of the nucleotide are indicated. Cryo-EM density is shown as a meshwork. The protein is shown in ribbon representation, nucleotide in stick representation and Mg^2+^ ion as a green sphere. (B to D) Zoomed-in views of pore-loops PL1 (B), PL2 (C) and PL3 (D) in relationship to the bound N-terminal peptide of RbcL-A. Peptide and side chains shown in stick representation. (E) Hydrogen bonds formed by the staggered pore-loops PL1 of Rca3 to Rca6 (Pro69) with the backbone of the RbcL N-terminal peptide at residues Lys11, Gln9, Lys7 and Gln5, respectively. (F) Cryo-EM density of the aromatic residues Tyr116 and Phe110 in pore-loops PL2 of Rca1 to Rca6.

**Figure S6.**
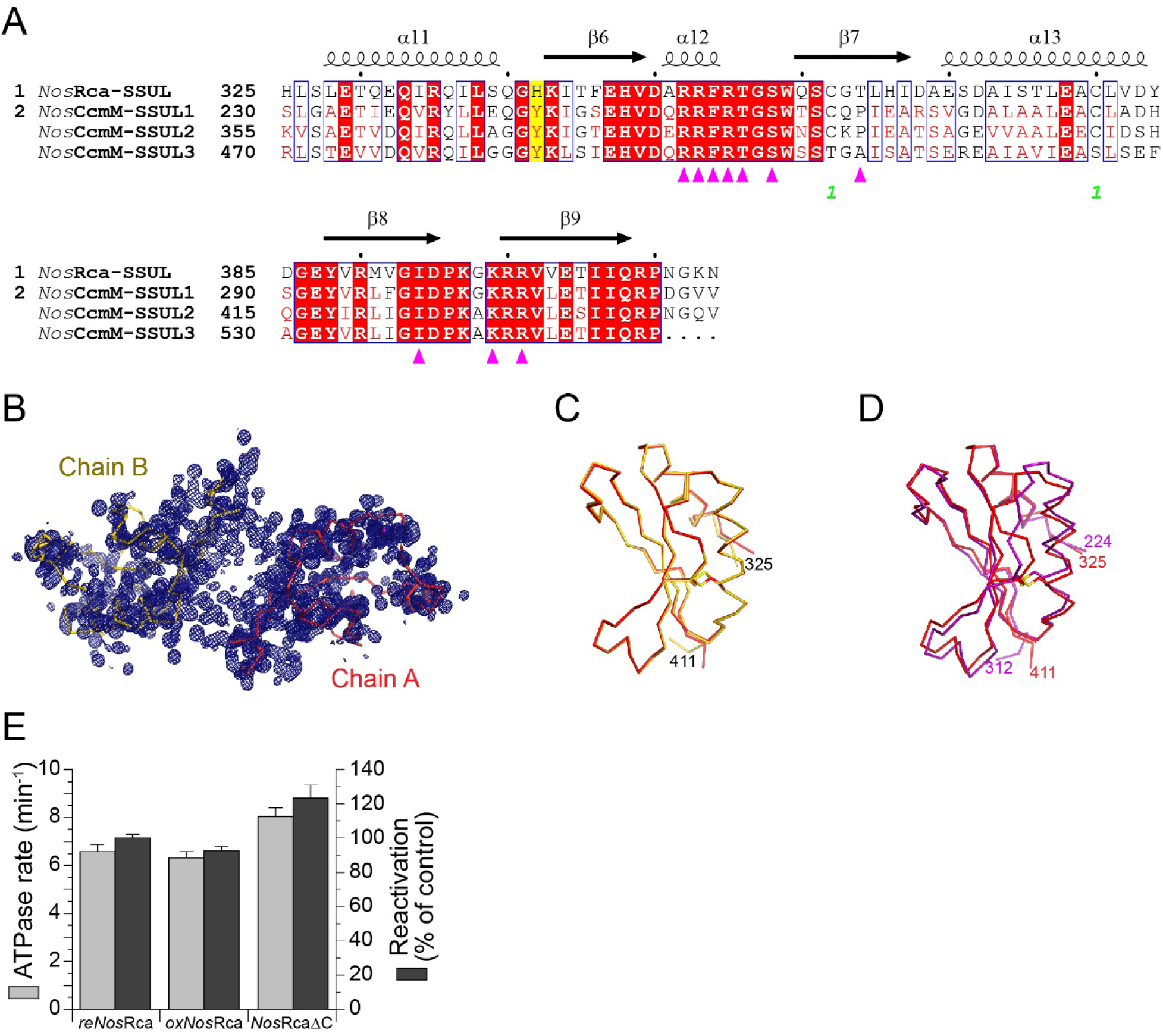
Role of the SSUL Domains of *Nos*Rca in Rubisco Binding. (A) Sequence alignment of the SSUL domain of *Nos*Rca (*Nos*SSUL) and the three SSUL modules of the scaffolding protein CcmM from *Nostoc* sp. PCC 7120. Secondary structure elements for *Nos*SSUL are indicated above the sequence. Similar residues are shown in red and identical residues in white on a red background. Blue frames indicate homologous regions. Yellow background indicates systematic differences between *Nos*SSUL and the three SSUL modules of *Nos*CcmM. The Uniprot accession codes for the sequences are: P58555, *Nos*SSUL domain from *Nostoc* sp. PCC 7120; Q8YYI3, CcmM from *Nostoc* sp. PCC 7120. (B) Unbiased experimental electron density for the asymmetric unit in crystals of the *Nos*SSUL domain at 1.4 Å resolution. The density after MAD phasing and density modification contoured at 1.5 σ is shown as a blue meshwork. The asymmetric unit contained two nearly identical copies (r.m.s.d. 0.43 Å for all Cα positions) (for data collection and refinement statistics, see Table S2). Chains A and B of the *Nos*SSUL domain are shown as Cα traces in red and yellow, respectively. (C) Superposition of chain A with chain B in the asymmetric unit of the *Nos*SSUL crystal lattice. The r.m.s.d. for matching Cα positions is 0.27 Å. Chain termini are indicated. (D) Superposition of the *Nos*SSUL domain with the SSUL1 domain of CcmM from *Synechococcus elongatus* PCC 7942 (*Se*SSUL; PDB: 6HBA). The *Se*SSUL is shown in purple. The r.m.s.d. for matching Cα positions is 1.0 Å (82 matching Cα positions). (E) ATPase and reactivation rates of reduced (re*Nos*Rca), oxidized (ox*Nos*Rca) and C-terminally truncated (*Nos*RcaΔC) *Nos*Rca. ATPase rates were measured in the absence of Rubisco at 20 mM KCl (see STAR Methods). CO_2_ fixation was measured for 8 min and set to 100 % for re*Nos*Rca. Error bars represent SD of at least three independent experiments.

**Figure S7.**
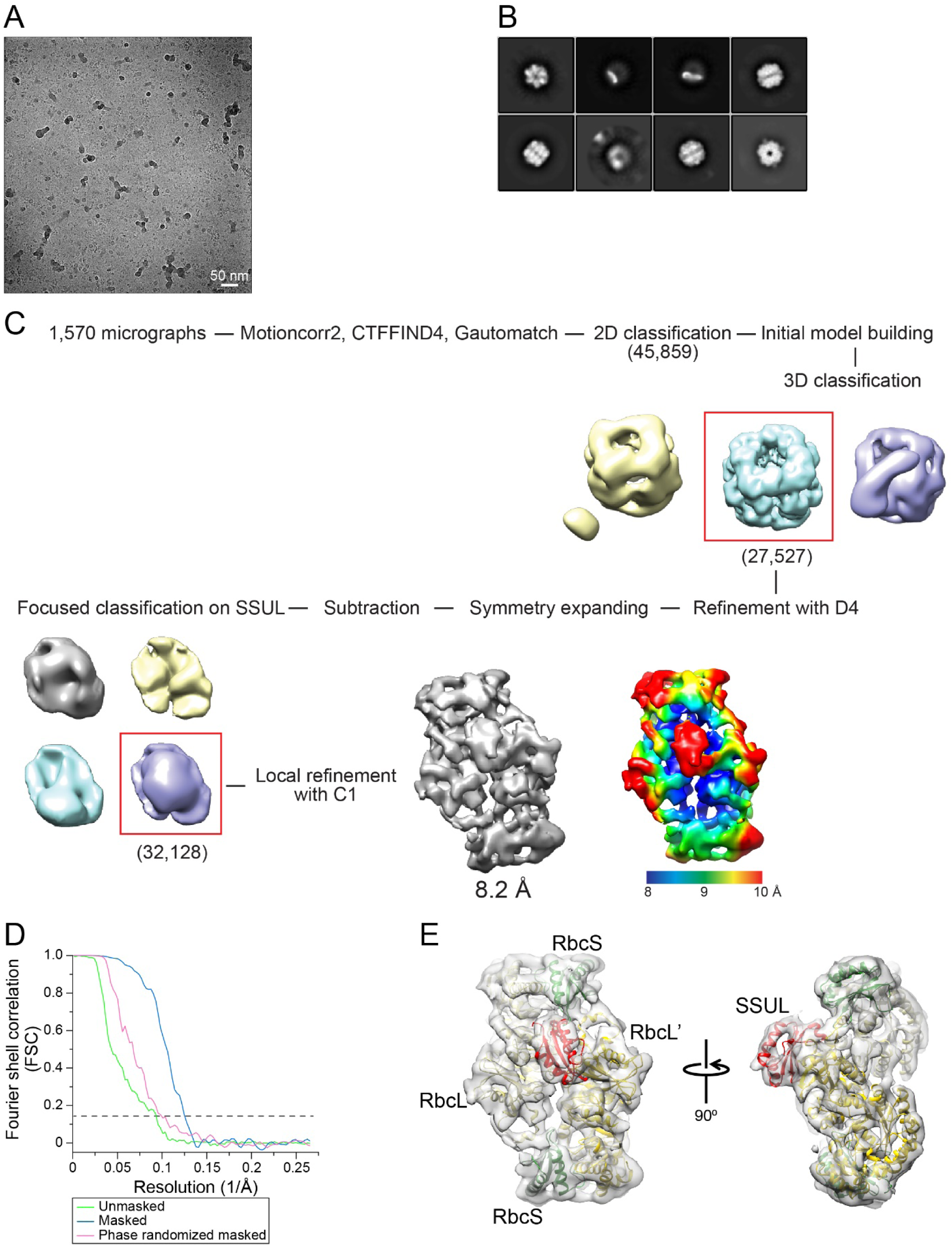
Cryo-EM Single-particle Reconstruction of NosRca:Rubisco Complex. (A) A representative micrograph of *Nos*Rca:Rubisco complexes. (B) 2D class averages of complexes in (A). (C) The single-particle data processing workflow for *Nos*Rca:Rubisco complex and local resolution map. The color gradient from blue to red indicates local resolution from 8.0 to 10 Å. (D) Gold-standard FSC curves of the *Nos*Rca:Rubisco reconstruction. The resolution is ∼8.2 Å at the FSC cutoff of 0.143 for the masked and B-factor sharpened curve. (E) Density map of 3D reconstruction from cryo-EM and single-particle analysis of 2RbcL-2RbcS-SSUL units at ∼8.2 Å resolution. The structural model in ribbon representation is docked into the cryo-EM density. The RbcL subunits in gold, RbcS in green and the bound SSUL domain in red.

## SUPPLEMENTAL TABLES

**Table S1.** Molar Mass and Hydrodynamic Radius of Proteins Determined by SEC-MALS.

| Protein | Measured Molar mass (Da) | Theoretical MW (Da) |  | Hydrodynamic radius (nm) |
| --- | --- | --- | --- | --- |
|  |  | monomer | hexamer |  |
| *NosRca*WT | ~274000 | 46668.03 | 280008.18 | ~6.2 |
| *NosRca*WT/ATP | ~255200 |  |  | ~5.7 |
| *NosRca*WT/ATPγS | ~255800 | | | ~6.0 |
| *NosRca*WT (L244D/N248A/L250A) | ~279300 | 46453.66 | 278721.96 | ~6.0 |
| *NosRca*WT (V91E/R92G/G93N/L244D/N248A/L250A) | ~267400 | 46441.56 | 278649.36 | ~5.9 |
| *NosRca*ΔC | ~199700 | 32865.57 | 197193.42 | ~5.4 |
| *NosRca*ΔC/ATP | ~212600 | | | ~5.5 |
| *NosRca*ΔC/ATPγS | ~196200 | | | ~5.0 |
| *NosRca*ΔC (V91E/R92G/G93N) | ~195000 | 32722.28 | 196333.68 | ~5.3 |
| *NosRca*ΔC (P140G/Y143A/D144A) | ~197000 | 32558.21 | 195349.26 | ~5.0 |
| *NosRca*ΔC (L244D/N248A/L250A) | ~197400 | 32651.2 | 195907.2 | ~5.1 |
| *NosRca*-SSUL | ~10430 | 10938 |  | ~1.7 |

**Table S2.**
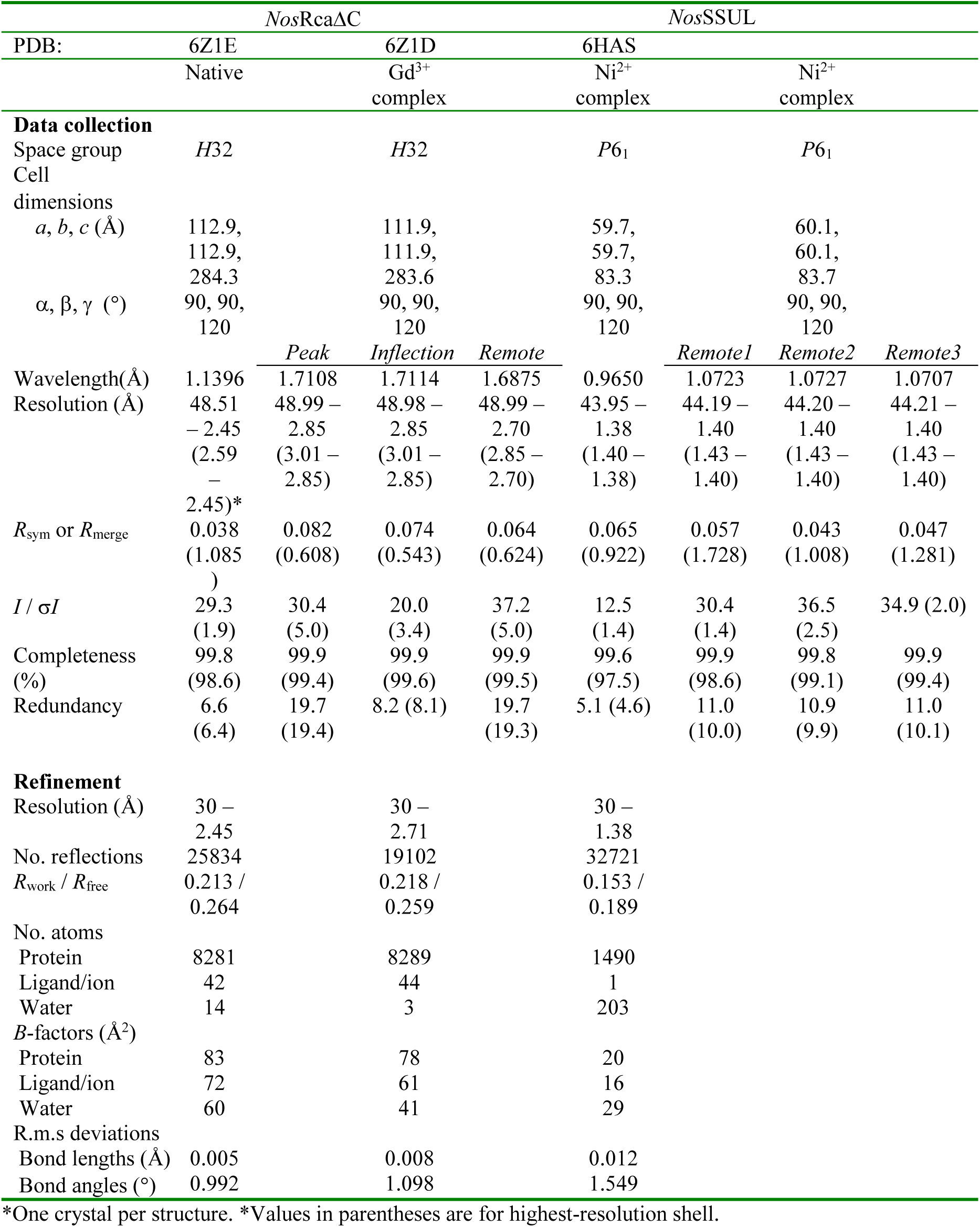
Data Collection, Phasing and Refinement statistics.

**Table S3.** Cryo-EM Statistics and Model Validation.

|  | *NosRca*□*C:Rubisco*<br>(EMDB: EMD-11028;<br>PDB: 6Z1F) | *NosRca:Rubisco*<br>(EMDB: EMD-11029;<br>PDB: 6Z1G) |
| --- | --- | --- |
| **Data collection and processing** |  |  |
| Magnification | 105000 | 22000 |
| Voltage (kV) | 300 | 200 |
| Electron exposure (e<sup>-</sup>/Å<sup>2</sup>) | 60 | 47 |
| Defocus range (μm) | -0.7 to -2.5 | -1 to -4.5 |
| Pixel size (Å) | 0.8512 | 1.885 |
| Symmetry imposed | C1 | C1 |
| Initial particle images (no.) | 335,474 | 298,336 |
| Final particle images (no.) | 21,149 | 32,128 |
| Map resolution (Å) | 2.86 | 8.2 |
| FSC threshold | 0.143 | 0.143 |
| Map resolution range (Å) | 2.4-23.57 | 8.0-50.0 |
| **Refinement** |  |  |
| Initial model used (PDB code) | 6Z1E, 1UZH |  |
| Model resolution (Å) | 3.2 | 8.8 |
| FSC threshold | 0.5 | 0.5 |
| Model resolution range (Å) | 2.7-40 | 6.9-130 |
| Map sharpening B factor (Å<sup>2</sup>) | -7.6 | -200 |
| **Model composition** |  |  |
| Non-hydrogen atoms | 49823 | 9262 |
| Protein residues | 6255 | 1168 |
| Ligands | MG: 12<br>CAP: 7<br>AGS: 5<br>ADP: 1 |  |
| B factors (Å<sup>2</sup>) |  |  |
| Protein | 65.1 | 285.1 |
| Ligand | 65.4 |  |
| **R.m.s. deviations** |  |  |
| Bond lengths (Å) | 0.008 (57) | 0.013(0) |
| Bond angles (°) | 1.172 (56) | 1.668(1) |
| **Validation** |  |  |
| MolProbity score | 1.33 | 2.25 |
| Clashscore | 2.79 | 5.98 |
| Poor rotamers (%) | 1.00 | 5.65 |
| **Ramachandran plot** |  |  |
| Favored (%) | 96.17 | 94.89 |
| Allowed (%) | 3.72 | 5.65 |
| Disallowed (%) | 0.11 | 0.43 |

## SUPPLEMENTAL DATA

**Data S1.**
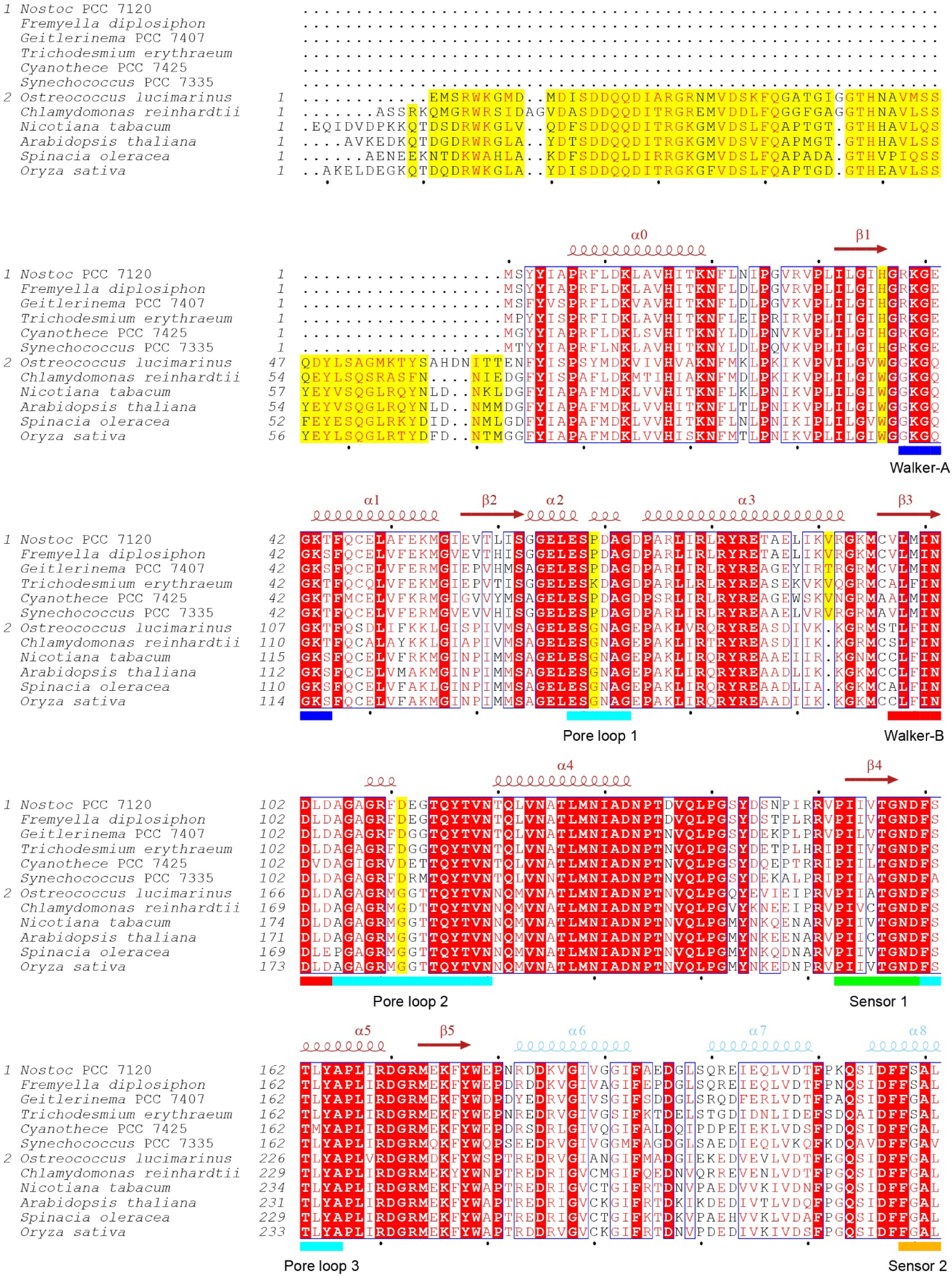
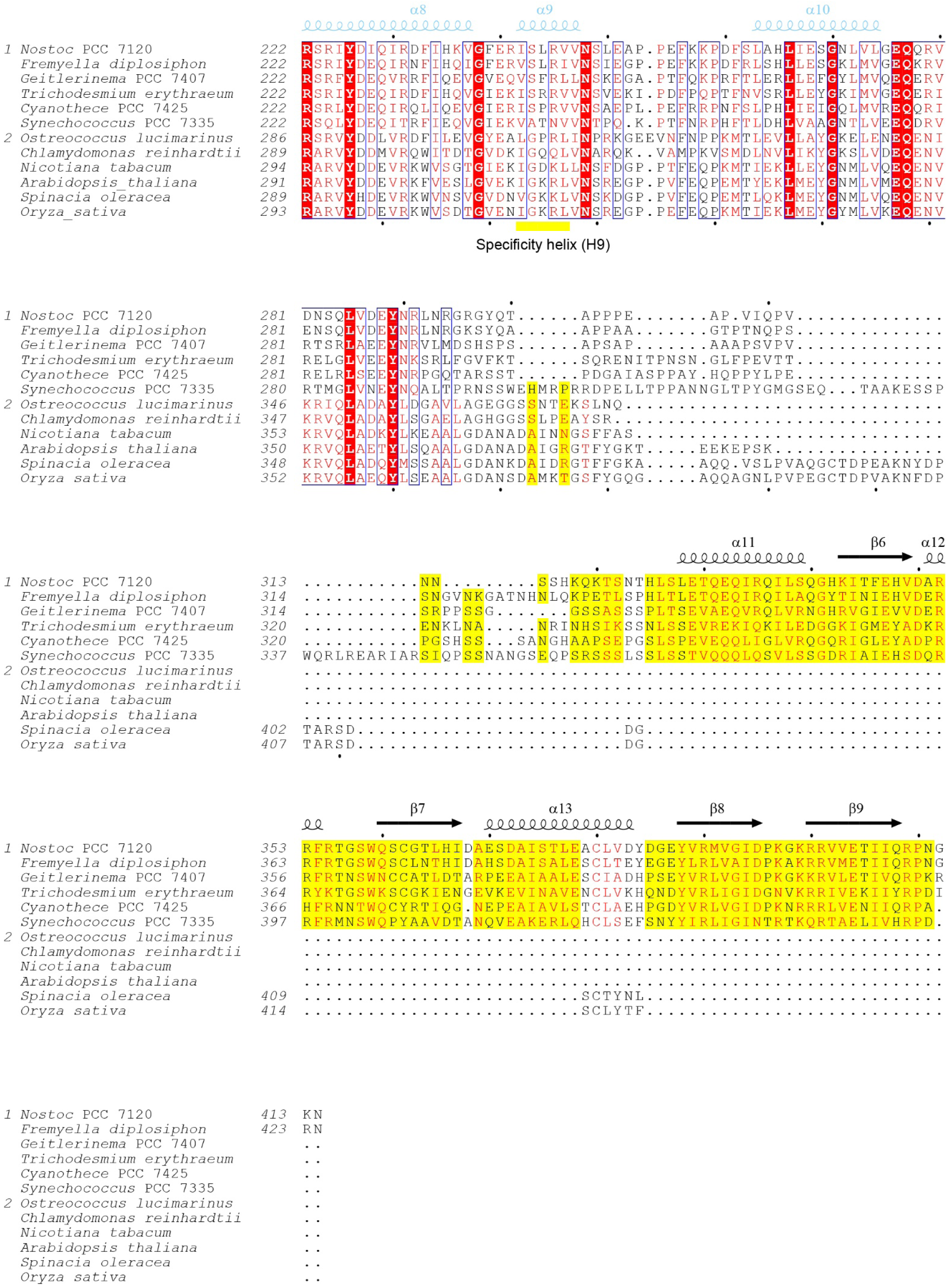
Alignment of Cyanobacterial ‘Rca-like’ and Plant Rca Sequences. Amino acid sequences of representative sets of cyanobacterial ‘Rca-like’ (group 1) and eukaryotic green lineage Rca proteins (group 2) were aligned using the EBI Clustal-O server (https://www.ebi.ac.uk/Tools/msa/clustalo/). Secondary structure elements for Rca from *Nostoc* sp. PCC 7120 are indicated above the sequences. Similar residues are shown in red and identical residues in white on a red background. Blue frames indicate homologous regions. Yellow background indicates systematic differences between cyanobacterial and eukaryotic sequences. Canonical elements of AAA+ domains are indicated by colored bars: Dark blue, Walker-A motif; red, Walker-B motif; green, sensor 1; orange, sensor 2. Teal bars indicate pore-loops PL1, PL2 and PL3. The yellow bar denotes the specificity helix H9. The Uniprot and Genebank accession codes for the sequences are: P58555, *Nostoc* sp. PCC7120; NZ JH930359.1, *Fremyella diplosiphon*; K9SEV2, *Geitlerinema* PCC7407; Q115H0, *Trichodesmium erythraeum*; B8HYF8, *Cyanothece* PCC7425; B4WNZ5, *Synechococcus* PCC7335; D8TZU3, *Volvox carteri*; A4RW20, *Ostreococcus lucimarinus*; Q6SA05, *Chlamydomonas reinhardtii*; Q40460, *Nicotiana tabacum*; P10896, *Arabidopsis thaliana*; P10871, *Spinacia oleracea*; P93431, *Oryza sativa*. The Figure was generated with ESPript (Gouet et al., 1999).

**Data S2.**
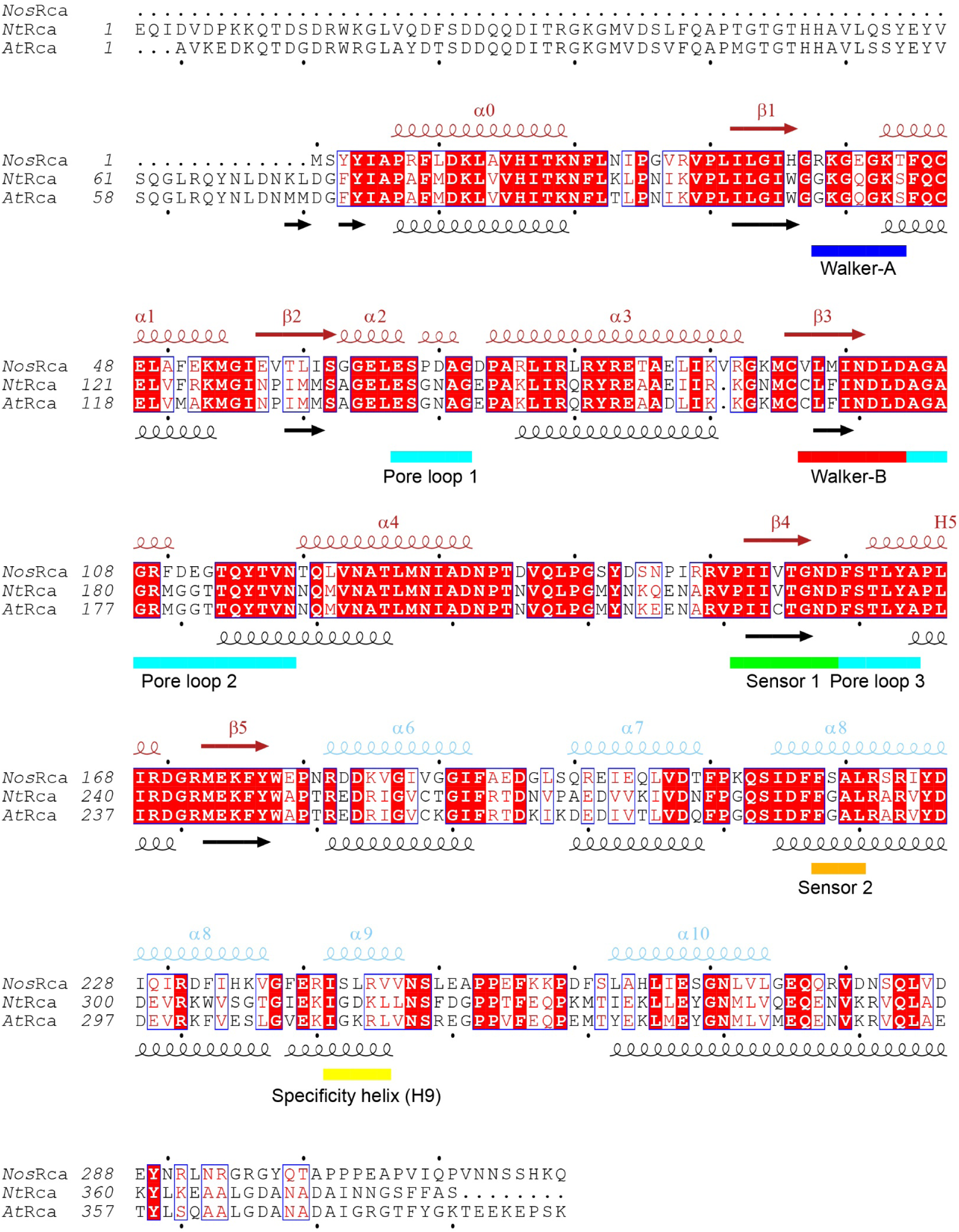
Sequence Alignment of *Nos*Rca, *Nt*Rca and *At*Rca. Amino acid sequences of *Nos*Rca, *Nt*Rca and *At*Rca were aligned using the EBI Clustal-O server (https://www.ebi.ac.uk/Tools/msa/clustalo/). Secondary structure elements for *Nos*Rca and plant Rca proteins are indicated above and below the sequences, respectively. Similar residues are shown in red and identical residues in white on a red background. Blue frames indicate homologous regions. Canonical elements of AAA+ domains are indicated by colored bars: Dark blue, Walker-A motif; red, Walker-B motif; green, sensor 1; orange, sensor 2. Teal bars indicate pore-loops PL1, PL2 and PL3. The yellow bar denotes the specificity helix H9. The Uniprot accession codes for the sequences are: P58555, *Nostoc* sp. PCC 7120; Q40460, *Nicotiana tabacum* and P10896, *Arabidopsis thaliana*.

